# Emergence of multiphase condensates from a limited set of chemical building blocks

**DOI:** 10.1101/2023.11.30.569439

**Authors:** Fan Chen, William M. Jacobs

## Abstract

Biomolecules composed of a limited set of chemical building blocks can co-localize into distinct, spatially segregated compartments known as biomolecular condensates. Although recent studies of intracellular condensates have shown that coexisting, immiscible condensates can form spontaneously via phase separation, it has remained unclear how coexisting and multiphase condensates assemble from chemical building blocks with limited specificity. Here we establish a connection between the interdependencies among biomolecular interactions and the thermodynamic stability of multiphase condensates. We then introduce an inverse design approach for computing the minimum interaction specificity required to assemble condensates with prescribed molecular compositions in a multicomponent biomolecular mixture. As a proof of principle, we apply our theory to design mixtures of model heteropolymers using a minimal number of distinct monomer types, and we use molecular simulations to verify that our designs produce coexisting condensates with the target molecular compositions. Our theoretical approach explains how multiphase condensates arise in naturally occurring biomolecular mixtures and provides a rational algorithm for engineering complex artificial condensates from simple chemical building blocks.

## I. INTRODUCTION

Biomolecules such as proteins and nucleic acids can spontaneously co-localize to form multicomponent condensates via the thermodynamically driven process of phase separation [1–4]. A wide variety of different condensates have been observed in living cells, each of which is associated with a distinct macromolecular composition and the ability to recruit specific client molecules, including enzymes and metabolites [5–7]. Moreover, condensate assembly and disassembly can be regulated by small changes in macromolecular concentrations and the chemical state of the intracellular environment, allowing for dynamic responses of biomolecular co-localization to external stimuli [8–13]. Condensate assembly therefore represents an important mechanism of self-organization in cellular biology, and considerable effort has been devoted to identifying so-called “grammars” that connect the molecular-level properties of specific biomolecules to the condensates that they tend to form [14–18].

Nonetheless, explanations for the observed diversity of condensates, which assemble in a controlled fashion in an extremely heterogeneous intracellular environment, have been elusive. The fundamental paradox stems from the well established biophysical properties of condensate-forming biomolecules. Because key components of condensates, such as intrinsically disordered proteins and many RNAs, are conformationally heterogeneous, these molecules tend to engage in relatively promiscuous, “multivalent” interactions in condensed phases that are fluid in nature [14, 19–22]. Furthermore, many of the molecular grammars that have been developed to describe individual systems, such as arginine/tyrosine compositions of FUS condensates [16] or the aromatic composition of prion-like domain condensates [17], are common to a wide variety of intracellular biomolecules which do not necessarily co-localize into the same condensates. It has thus been unclear how the *compositional specificity* [23, 24] of multicomponent condensates arises from relatively disordered macromolecules, non-stoichiometric interactions, and existing theories of molecular grammars [25]. Resolving this question is essential to understanding how phase separation can give rise to complex intracellular spatial organization [25].

We propose that this question can be resolved by focusing on the *interaction specificity* of biomolecules. More precisely, we ask what degree of interaction specificity is required to achieve the observed spatial organization in a multicomponent system. Theoretical predictions can then be tested either by quantifying the interaction specificity in a mixture of biomolecules that is observed to phase-separate into distinct coexisting condensates, or by designing “minimal-complexity” biomolecular mixtures—i.e., mixtures that possess the minimum required interaction specificity—that phase-separate into coexisting condensates with prescribed molecular compositions. This approach builds on recent theoretical progress in modeling multicomponent phase separation [25– which aims to predict multiphase coexistence using simplified descriptions of biomolecular interactions. However, by focusing on interaction specificity, we can establish a direct connection between multiphase coexistence and the required *physicochemical properties* of biomolecular mixtures.

In this article, we develop a general theoretical description of biomolecular interaction specificity and show how it relates to the thermodynamic stability of coexisting multicomponent condensates. We then introduce an “inverse design” approach [27, 28] for computing the minimum interaction specificity required to form immiscible condensates via phase separation. In addition, our approach establishes a straightforward algorithm for designing minimal-complexity mixtures of biomolecules or synthetic macromolecules, subject to system-specific physicochemical constraints, that phase-separate into coexisting phases with prescribed molecular compositions. We demonstrate how this algorithm can be used in practice by designing mixtures of heteropolymers that phase-separate into non-trivial coexisting phases while utilizing a surprisingly small number of distinct monomer types. Finally, we perform molecular dynamics simulations of phase coexistence with these heteropolymer mixtures to validate our theory and inverse design approach. Our results suggest a unifying framework for exploring molecular grammars in biomolecular systems, as well as a practical strategy for engineering complex artificial condensates using experimentally realizable molecules.

**FIG. 1.**
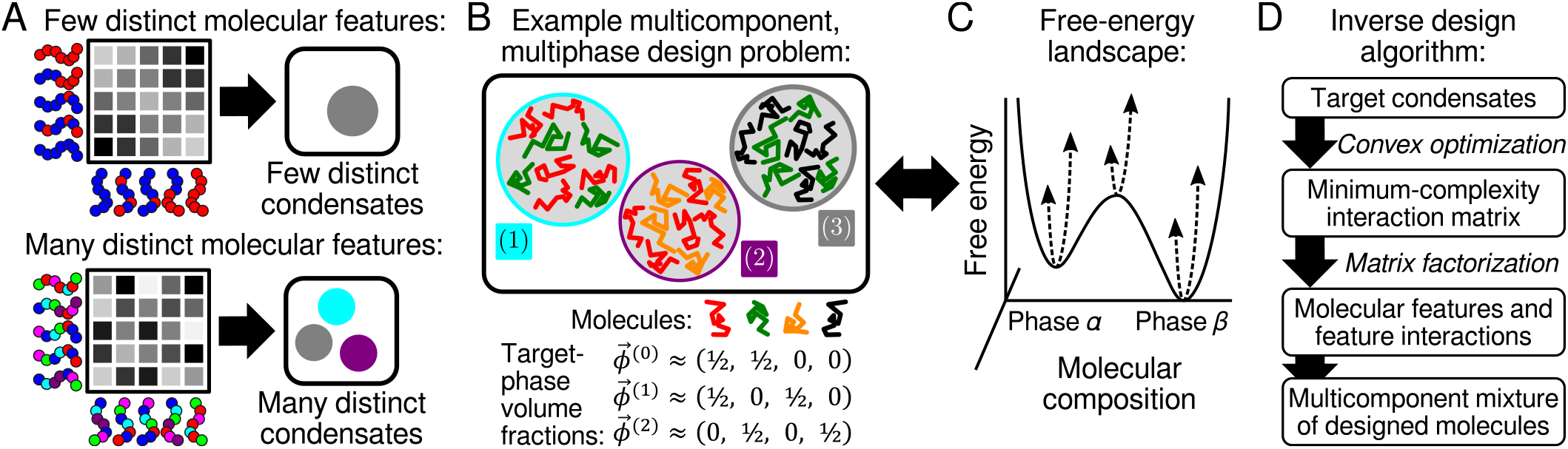
Biomolecular interaction interdependencies and inverse design approach. (A) The interactions between biomolecules in a multicomponent mixture can be described by a pairwise interaction matrix. In a system with few molecular features, such as polymers with a small number of distinct monomer types, the elements of the interaction matrix tend to be interdependent (top). By contrast, in a mixture with many distinct molecular features, the interaction-matrix elements can be independently tuned (bottom). (B) An example multiphase design problem. Here, four molecular species (red, green, gold, and black) phase-separate into three immiscible condensates with prescribed molecular volume fractions 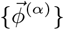. (C) A multiphase design problem corresponds to a high-dimensional free-energy landscape, where each target phase (such as the two shown here) specifies a local minimum. (D) To solve the design problem, we first use convex optimization to find the *minimum-complexity* interaction matrix, requiring the smallest number of distinct molecular features, that produces the desired free-energy landscape. We then factorize the minimum-complexity interaction matrix to construct a multicomponent mixture of designed molecules.

## II. RESULTS

### A. Describing interdependencies among biomolecular interactions

To develop a theoretical description of biomolecular interaction specificity, we adopt a coarse-grained representation of biomolecules in a multicomponent fluid. Specifically, we assume that *pairwise* interactions can be used to describe the net attraction or repulsion between molecular species [25]. This approach implies that the chemical potential *µ*_*i*_ of each macromolecular species *i* = 1, …, *N* depends linearly on the symmetric pairwise interaction matrix *ϵ*,

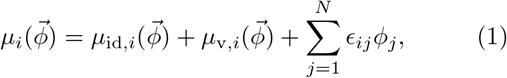

where *ϕ*_*i*_ represents the molecular volume fraction (i.e., the concentration times the molecular volume, *v*_*i*_) of each molecular species *i*. The first two terms of Eq. (1) account for the ideal, 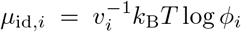, and steric, 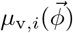, contributions to the chemical potential, which are both independent of *ϵ*. Eq. (1) provides a coarsegrained description of any homogeneous fluid [37]. However, we make the key approximation that *ϵ* is independent of 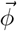; this means that the net interaction *ϵ*_*ij*_ between molecules of types *i* and *j* does not depend on the presence of other components in the mixture and is thus the same in all phases. This approximation underlies the Flory–Huggins [38], regular solution [39], and van der Waals models [37], and is consistent with field-theoretic treatments of heteropolymer mixtures [40], with appropriate choices of 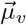. Later on, we will use molecular simulations to show that this approximation can yield a remarkably accurate description of multicomponent heteropolymer solutions.

The pairwise interaction matrix provides a systematic way of describing the interdependencies among biomolecular interactions in a multicomponent mixture. In a full-rank interaction matrix, there are *N*(*N* + 1)*/*2 independent pair interactions. However, because biomolecular interactions are typically governed by a limited set of physicochemical features, it is conceivable that not all pair interactions are independent of one another. For example, in a mixture of molecules that interact via hydrophobic forces alone, there might be only *N* independent parameters—namely, the hydrophobicity of each molecular species—that determine the interaction matrix. In general, we expect that various modes of interaction, including electrostatic, cation-*π*, and hydrogen bonding, among others [14], contribute to the net interactions among biomolecules in a multicomponent mixture. We formalize this concept of interaction interdependency (Fig. 1A) by factorizing *ϵ* in terms of a molecular feature matrix, *W* ∈ *ℛ*^*N ×r*^, and a feature interaction matrix, *u* ∈ *ℛ*^*r×r*^ [34],

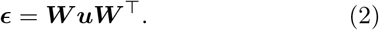

Here, a “feature” refers to a linearly independent molecular property, which could represent a literal molecular building block such as a nucleotide or an amino acid, or alternatively an emergent property such as a sequence motif or pattern. Each row vector of *W* describes the appearance of these features in a particular molecular species, while the matrix *u* describes the interactions among all pairs of features. Importantly, the rank *r* indicates the number of distinct features that are available, and thus controls the number of elements of *ϵ* that can be independently tuned. An interaction matrix can only be realized in a biomolecular system if *r* does not exceed the number of available features.

### B. Relating interaction interdependencies and biomolecular condensate thermodynamics

Next, to relate the number of available molecular features to the formation of coexisting biomolecular condensates, we take advantage of an *inverse design* approach to condensate thermodynamics [27, 28]. In this approach, we prescribe a set of *K* target condensed phases (i.e., condensates) with defined molecular compositions (Fig. 1B). Each target condensate *α* = 1, …, *K* is described by a vector of molecular volume fractions 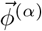. For these condensed phases to be thermodynamically stable, each 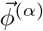 must correspond to a local minimum on a free-energy landscape 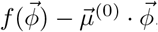, where 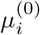 is the chemical potential of molecular species *i* in the dilute phase [37]. From the relationship between the free-energy density *f* and the chemical potentials, *µ*_*i*_ = ∂*f/*∂*ϕ*_*i*_, this stability condition implies that 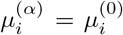 for all condensed phases and molecular species. The stability condition also implies that the Hessian of the free-energy density must be positive definite in each condensed phase *α*, 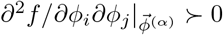.

An interaction matrix that solves this inverse design problem establishes a free-energy landscape, following the assumptions of Eq. (1), that is consistent with the prescribed *K* condensed-phase molecular compositions (Fig. 1C). As such, any mixture with overall macromolecular concentrations within the convex hull of the target-phase and dilute-phase concentrations 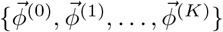 can phase-separate to form the prescribed condensed phases [28]. (Whether some or all of these phases form in practice depends on kinetic considerations, such as nucleation dynamics [25, 27], that are not the focus of the present work.) However, the interaction matrix that solves the design problem is typically not unique. In particular, there may be low-rank interaction matrices, for which *r < N*, that result in—or at least closely approximate—the prescribed condensed-phase molecular compositions. Our goal is to find such low-rank solutions, since they reduce the number of molecular features that are required to achieve the prescribed set of coexisting condensates.

To relate the minimum rank *r* to a target set of condensed phases 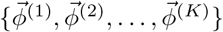, we consider an interaction matrix *ϵ* that solves this inverse problem. Perturbing *ϵ* by an error matrix Δ*ϵ* results, to lowest order, in a shift of the condensed-phase compositions (see SI),

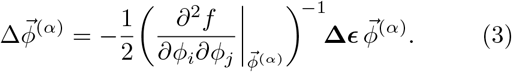

Eq. (3) says that the concentrations in the *α* phase are most sensitive to perturbations in the direction of the eigenvector, 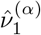, that corresponds to the minimum eigenvalue, 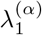, of 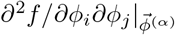. We can therefore relate the Frobenius norm of a random error matrix Δ*ϵ* to the relative change in the molecular compositions of the *α* phase in this least stable direction, 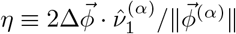 (see SI),

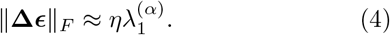

Eq. (4) is useful because it relates the interaction-matrix perturbation to its effect on the phase behavior in the worst-case scenario. Thus, to solve the inverse design problem to within a composition tolerance of *η*, we must limit deviations of the interaction matrix such that 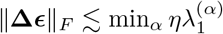.

We can now determine the minimum required interaction specificity by attempting to reduce the number of molecular features, *r*≤*N*, used to represent the interaction matrix *ϵ*. Doing so yields a low-rank approximation, *ϵ*_*r*_, for the *N×N* interaction matrix. Here we apply the Eckart–Young–Mirsky (EYM) theorem [41], which says that ∥Δ*ϵ*∥ _*F*_ = ∥*ϵ*_*r*_−*ϵ*∥_*F*_ is minimized by eliminating the *N*−*r* smallest singular values of *ϵ*. Accordingly, the minimum number of molecular features, *r*, that successfully solves the inverse design problem must satisfy

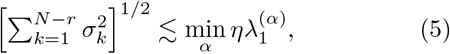

where *σ*_1_≤*σ*_2_≤ … ≤*σ*_*N*_ are the singular values of *ϵ*. Eq. (5), which relates the singular values of the interaction matrix *ϵ* to thermodynamic properties of the coexisting condensates, is the central result of our theoretical approach. This equation implies that the inverse design problem can be solved—using the smallest possible number of distinct molecular features—by optimizing *ϵ* to simultaneously minimize the smallest singular values and maximize the stabilities of the target condensed phases. In this way, our inverse design approach establishes a direct connection between the interdependencies of pairwise biomolecular interactions and the complexity of the phase-separated condensates that they can form, irrespective of specific details of the molecular features in a particular biomolecular system.

### C. Designing low-rank interaction matrices and minimal-complexity biomolecules

In practice, we implement this theory for designing low-rank interaction matrices using convex optimization [42] (Fig. 1D). Then, after the required number of molecular features (i.e., the pairwise interaction matrix rank *r*) has been determined, we design molecular mixtures by factorizing *ϵ*, subject to appropriate physicochemical constraints. We discuss each step in turn.

#### Step 1: Optimization of component-wise interactions

First, we define a convex relaxation of the inverse design problem, such as the example condensate design problem illustrated in Fig. 1B, following the strategy introduced in Refs. [27] and [28]. This step introduces controlled approximations to transform the nonlinear thermodynamic stability and design constraints described in Sec. II B into a convex optimization problem (see SI for complete details). This convex relaxation is defined by the target condensed-phase volume fractions 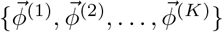, the molecular volumes of the *N* molecular species, and a minimum stability criterion *λ*_min_, which imposes a lower bound on 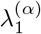 for each condensed phase *α*. By satisfying the constraints of this convex optimization problem, we identify a “solution space” of pairwise interaction matrices that closely approximate the target free-energy landscape. Importantly, convexity implies that this solution space can be efficiently computed, or otherwise proven to be infeasible if solutions to the inverse design problem do not exist [42–44].

Within this solution space of pairwise interaction matrices, we wish to find a matrix with the fewest required number of molecular features, in accordance with Eq. (5). We therefore minimize the nuclear norm of the interaction matrix, 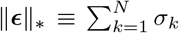, which is a convex relaxation of the matrix rank [45], subject to the aforementioned convex constraints. Including this objective function tends to reduce the magnitudes of the smallest singular values of *ϵ*. Moreover, minimizing ∥*ϵ*∥_*\**_ guarantees that the solution to the convex optimization problem is unique. We can then determine an upper bound on the minimum required number of features, *r*, using Eq. (5), and finally construct a rank-*r* approximation of the interaction matrix, *ϵ*_*r*_, by applying the EYM theorem.

#### Step 2: Factorization into molecular features

Second, having established the low-rank interaction matrix *ϵ*_*r*_, we find a molecular realization of the pairwise interactions by factorizing *ϵ*_*r*_ according to Eq. (2). The rank of the feature interaction matrix *u* must be exactly equal to *r*, so that exactly *r* features are required for each row vector of the molecular feature matrix *W*. An optimal factorization is obtained by minimizing the Frobenius norm of the reconstruction error,

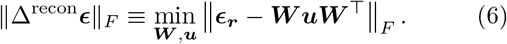

Up until this point, the design approach has been agnostic to the details of the molecular features. However, the optimal factorization given by Eq. (6) must typically satisfy additional constraints. Most importantly, a physically interpretable molecular feature matrix *W* typically has nonnegative entries, since each matrix element indicates the existence and magnitude of a distinct feature within a particular molecule. In this case, nonnegative matrix factorization (NMF) [46] must be used in place of eigenvalue decomposition when solving Eq. (6) (see SI). Further constraints may also be required depending on the specific requirements of the molecular system and the nature of the molecular features. For example, if each molecular feature represents the count of an amino acid type in a disordered polypeptide, then the elements of *W* must be nonnegative integers. Similarly, the feature interaction matrix *u* may either be fixed, as in the case of amino-acid interactions, or designable, as in the case of nucleic acid motifs [47]. As a concrete example, a specific heteropolymer model will be considered in detail below. Regardless of these system-specific constraints, the reconstruction error must obey 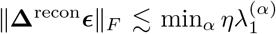 in order for the designed molecular mixture to form the prescribed condensed phases.

In summary, our inverse design approach establishes an upper bound on the number of distinct molecular features that are required to form multiple coexisting condensates with prescribed molecular compositions. We can then devise a specific molecular implementation, using the minimal number of molecular features, via a second optimization step. Next, we will demonstrate this two-step optimization procedure using a prototypical model of phase-separating polymers.

### D. Designing phase-separating heteropolymers using a minimal number of monomer types

To demonstrate the validity of our theory (Sec. II B) and design approach (Sec. II C), we consider a common simulation model of sequence-dependent heteropolymers. In this model, polymer species *i* = 1, …, *N* are linear chains of *L*_*i*_ monomers, which are chosen from a library of *r* monomer types (see SI for model details). Non-bonded monomers of types *a, b* = 1, …, *r* interact via a cut-and-shifted Lennard-Jones (LJ) pair potential [48] with monomer diameter *d*. Bonded monomers interact via finite-extensible nonlinear elastic (FENE) bonds [49]. Our aim is to design the sequences of polymers with maximum degree of polymerization *L*_max_, along with the nonpositive monomer interaction matrix *u*^LJ^, in order to form a set of *K* coexisting condensates with prescribed polymer compositions.

As an illustrative example, we consider the 4-component, 3-condensed-phase mixture depicted in Fig. 1B, which represents a nontrivial design problem due to the “enrichment” (i.e., high target volume fractions) of chains A and B in two distinct, immiscible condensed phases. Because we are designing linear polymers, we utilize the Flory–Huggins expression for the steric term in Eq. (1), 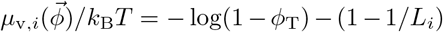, where 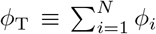 (see SI). In what follows, we choose the maximum degree of polymerization to be *L*_max_ = 20. By solving the minimum-nuclear-norm convex optimization problem as a function of the minimum thermodynamic stability criterion, *λ*_min_ (Sec. II C), we obtain the singular values of the optimized pair interaction matrix, *ϵ*, for this design problem (Fig. 2A). The nuclear norm of the optimized *ϵ* tends to increase with *λ*_min_, up to the point at which the design problem becomes infeasible, since imposing a stricter constraint on the free-energy landscape tends to shrink the interaction-matrix solution space. Accordingly, the singular values of *ϵ* tend to increase with *λ*_min_. We then compare the singular values with the right-hand side of Eq. (5), assuming a composition tolerance of *η* = 20%. Since one singular value lies below the bound in Fig. 2A, Eq. (5) predicts that one singular value can be eliminated while still satisfying the design problem. Thus, only *r* = 3 molecular features are required for the construction of this particular heteropolymer mixture.

**FIG. 2.**
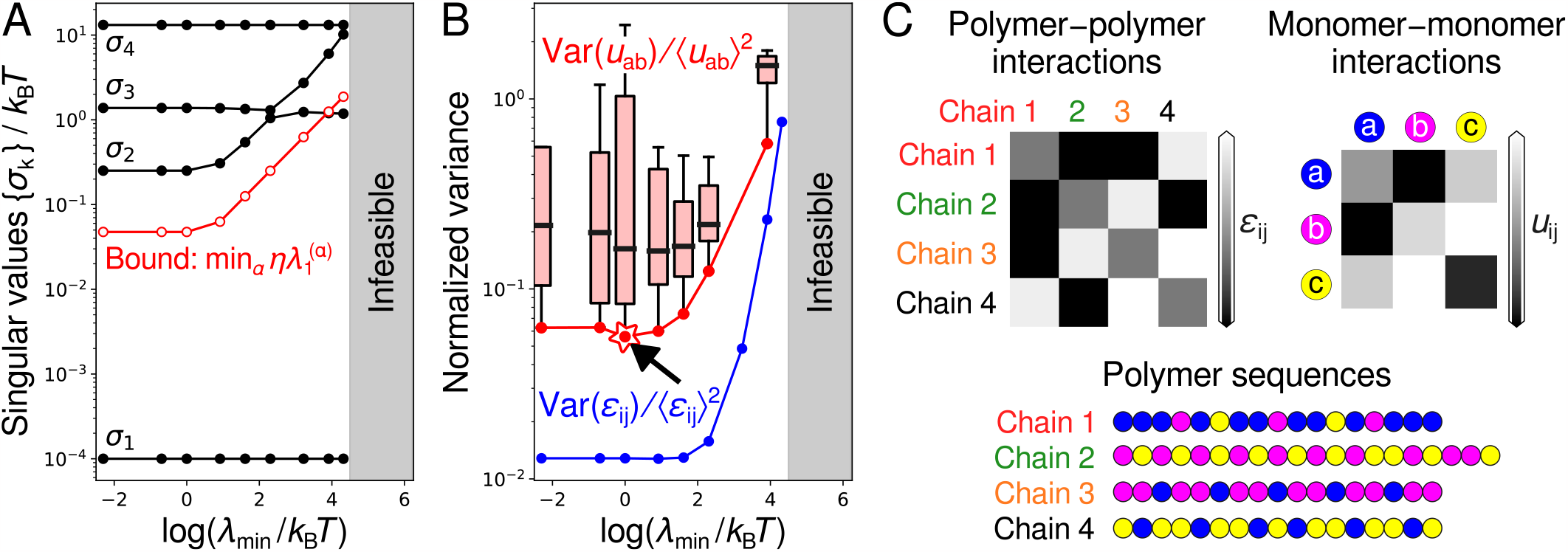
Optimization of minimal-complexity interaction matrices. (A) Inverse design of the 4-component multiphase system illustrated in Fig. 1B. The singular values of the optimized interaction matrix *ϵ*, obtained via nuclear-norm minimization (Sec. II C), as a function of the landscape stability criterion *λ*_min_. The singular values tend to increase as the condensed phases become more stable. Comparing the singular values with the bound, Eq. (5), indicates that the minimum interaction matrix rank is *r* = 3. (B) The normalized pairwise interaction variance, Var(*ϵ*_*ij*_)*/* ⟨*ϵ*_*ij*_ ⟩^2^, tends to increase with *λ*_min_. Box plots show the distribution of normalized monomer–monomer interaction variances obtained by nonnegative matrix factorization (NMF) of the optimized rank-3 polymer–polymer interaction matrices found in A, while treating the distinct monomer types as the molecular features. All NMF solutions satisfy 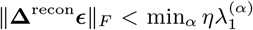. The solid red line indicates the Pareto front, representing the NMF solutions that simultaneously minimize Var(*u*_*ab*_)*/* ⟨*u*_*ab*_⟩^2^ and maximize 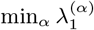. (C) The design solution corresponding to the starred location on the Pareto front (see arrow) in B. The polymer sequences consist of exactly *r* = 3 monomer types.

We now assume that the molecular features represent distinct monomer types in the LJ heteropolymer model. In this case, each row of the *N×r W* matrix is a count encoding of the number of occurrences of each monomer type in a heteropolymer sequence; consequently, all elements of *W* are nonnegative integers, and the row sums of *W* are bounded by the maximum degree of polymerization *L*_max_. Meanwhile, *u*^LJ^ is a nonpositive *r × r* interaction matrix determined by the attractive portion of the monomer–monomer LJ pair potential (see SI). Here, assuming that the molecular features represent distinct monomer types is equivalent to a pairwise-additive approximation for the polymer–polymer interactions, such that 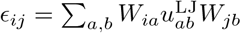. We emphasize that this approximation is not required by our theory, since we could instead factorize *ϵ* via an eigendecomposition of pairwise polymer–polymer interactions determined directly from simulations. However, we choose to make this assumption here to provide a transparent proof of principle. We note that this approximation establishes an upper bound on the required number of distinct monomer types, since the number of molecular features obtained from an eigen-decomposition of *ϵ* cannot be less than the number of distinct monomer types. In other words, while we can use this approximation to rigorously test our theory, the bound on the required number of monomer types could potentially be improved by relaxing this assumption.

Our next step is to solve Eq. (6) subject to the optimized low-rank interaction matrix, *ϵ*_*r*_, and the constraints on *W* and *u*. This mixed-integer quadratic non-negative matrix factorization (NMF) problem [46] can be solved using a stochastic optimization algorithm [50]. Specifically, we generate an ensemble of *W* matrices and iteratively solve for the interaction matrix *u* to ensure non positivity; we then utilize enhanced sampling to probe the tail of the reconstruction-error distribution to find the *W* matrix with the smallest ∥Δ^recon^*ϵ*∥_*F*_ (see SI). Representative results of this stochastic algorithm are shown in Fig. 2B. We find that the relative variance of the optimized monomer–monomer interactions, Var(*u*_*ab*_)*/* ⟨*u*_*ab*_⟩^2^, tends to increase with the minimum stability constraint *λ*_min_. This trend reflects the behavior of the polymer–polymer interaction matrix, *ϵ*_*r*_, that is being factorized, Var(*ϵ*_*ij*_)*/* ⟨*ϵ*_*ij*_⟩^2^, and suggests a fundamental trade-off: Designing highly stable condensates, which are least sensitive to interaction-matrix perturbations according to Eq. (3), requires highly dissimilar monomer– monomer interactions.

**FIG. 3.**
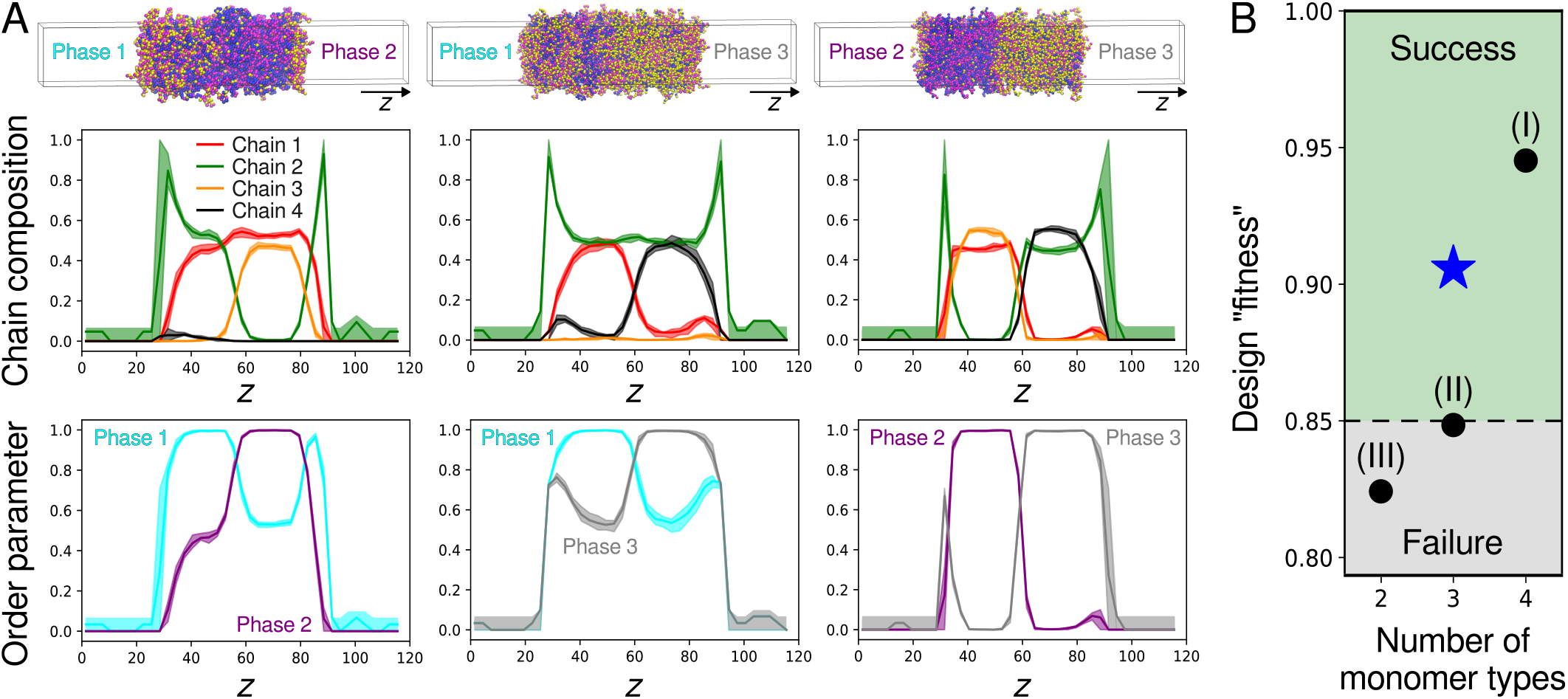
Validation of minimal-complexity heteropolymer designs via molecular simulation. (A) Direct-coexistence simulations of all pairs of condensed phases illustrated in Fig. 1B using the 3-monomer-type heteropolymer design shown in Fig. 2C. *Top:* Equilibrium configurations, color-coded by monomer type. *Middle:* Trajectory-averaged chain composition profiles along the direction of the simulation box perpendicular to the condensate interfaces. *Bottom:* Corresponding trajectory-averaged order parameter profiles, Eq. (7). Shaded regions indicate the statistical uncertainty associated with the mean profiles.(B) Comparison of the “fitness” metric (Sec. II E) for the heteropolymer design studied in A (blue star) and three alternative designs (black circles): (I) a 4-monomer-type design chosen to maximize the condensed-phase stabilities, (II) a 3-monomer-type design constructed by applying the EYM theorem to an interaction matrix that was *not* obtained via nuclear-norm minimization, and (III) a two-monomer-type design constructed by applying NMF with *r* = 2 (see SI for complete details of these alternative designs). Designs are classified as successes or failures based on a fitness threshold of 0.85 (see SI).

Finally, we use an optimized monomer-composition matrix, *W*, and monomer–monomer interaction matrix, *u*^LJ^, to construct a set of heteropolymer sequences for simulation. Because the pairwise-additive approximation relating *ϵ*_*ij*_ and 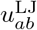 is most accurate when Var(*u*_*ab*_)*/* ⟨*u*_*ab*_⟩^2^ is small, we select monomer designs from the Pareto front shown in Fig. 2B. We then compose the polymer sequences from the monomer compositions by interleaving the different monomer types to minimize the blockiness of each sequence (see SI). An example outcome of this algorithm, which we will directly test via molecular simulation, is shown in Fig. 2C.

### E. Validating multicomponent, multiphase heteropolymer designs via molecular simulation

To test our LJ heteropolymer designs, we perform direct-coexistence molecular dynamics simulations [51] to examine the molecular compositions and immiscibility of the target condensates. In these simulations, all chain types are present in a constant-temperature, constant-volume simulation with periodic boundary conditions. However, the initial condition is chosen such that two different condensed phases, *α* ≠ *β*, are in contact with one another and also with a dilute phase. The condensed phases are thermodynamically stable if they remain phase-separated and immiscible once the simulation has relaxed to equilibrium; in practice, we ensure convergence by calculating simulation trajectories that are many times longer than the time required for all the chains in an *α* = *β* control simulation to mix completely (see SI). We carry out simulations for all pairs of condensed phases *α* ≠ *β* to confirm that they are mutually immiscible.

In Fig. 3A, we show equilibrium configurations of simulations and polymer composition profiles for the LJ heteropolymer design shown in Fig. 2C. We define an order parameter for each phase *α* based on the cosine similarity of the local polymer composition 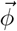,

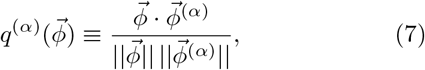

to distinguish the coexisting condensed phases (Fig. 3A). This order parameter is equal to one if the polymer composition of an equilibrated condensed phase matches that of target phase *α*. When coexisting condensed phases have “shared” components, meaning that one or more polymer species are enriched in multiple phases, the order parameters are not orthogonal, such that 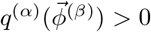. The order parameter profiles shown in Fig. 3A indicate that our 3-monomer-type LJ heteropolymer mixture indeed solves the design problem presented in Fig. 1B.

We can compare alternative heteropolymer designs by introducing a “fitness” metric computed from the order parameter profiles given by Eq. (7) (see SI). Using the fact that the minimum fitness of a valid solution is ≈0.85 (see SI), we can quantitatively assess whether each design is successful (Fig. 3B). The optimized 3-monomer-type design shown in Fig. 3A is classified as a success as expected, as is a 4-monomer-type design that has zero reconstruction error since the matrix factorization step, Eq. (6), is not required. By contrast, a 3-monomer-type design constructed by applying the EYM theorem to an interaction matrix that was *not* optimized to minimize ∥*ϵ*∥_*\**_, and thus does not satisfy Eq. (5), fails this test. Similarly, performing NMF to solve Eq. (6) using only two monomer types leads to a large reconstruction error that violates Eq. (5), and the simulated condensed phases are observed to mix. Although this empirical test does not preclude the possibility of a successful two-monomer-type design that manifests three molecular features via sequence-patterning effects, it is consistent with our prediction that three monomer types are required if the monomer types are assumed to be the features in the heteropolymer design step. (Detailed descriptions of these alternative designs and accompanying simulation data are provided in the SI.) Finally, we note that interesting interfacial effects, particularly between the condensed and dilute phases, are more prominent in the optimized 3-monomer-type design (Fig. 3A) than in the 4-monomer-type design. This behavior is not inconsistent with our theory or design approach, which focus solely on the molecular compositions of bulk phases, and likely reflects the reduced condensed-phase stabilities (and thus lowered surface tensions) of the 3-monomer-type design.

### F. Evaluating the pairwise approximation using simulations of successful condensate designs

Given the successful simulation test of our 3-monomer-type heteropolymer mixture, it is important to examine why our design approach works so well. Specifically, we can assess the pairwise-additive approximation for the polymer–polymer interactions, which is invoked when treating the monomer types as the molecular features, directly via simulation. We first consider the interactions between polymers in a dilute mixture. These interactions are quantified by computing the second virial coefficients between polymers, 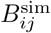, and compared to the second virial coefficients predicted by the Flory–Huggins model, 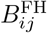. We find that 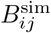 and 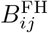 are positively correlated with a Pearson correlation coefficient of 0.67, which suggests non-negligible differences between the dilute-phase polymer–polymer interactions and the predictions of the pairwise-additive approximation (Fig. 4A).

**FIG. 4.**
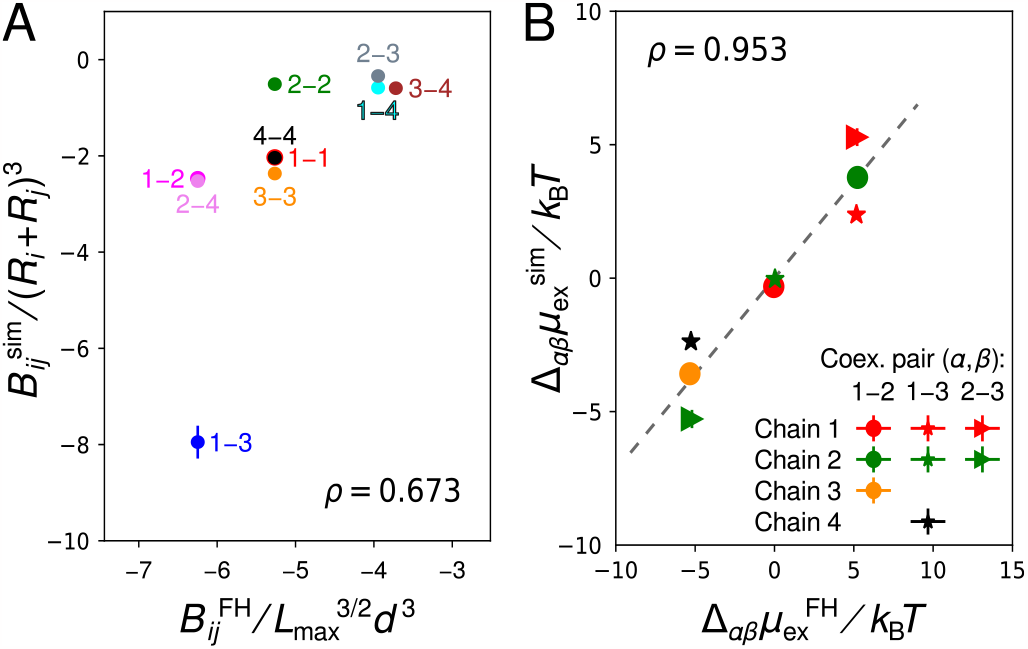
Validation of the pairwise-additive approximation for polymer–polymer interactions. (A) Correlation between the second virial coefficients obtained from simulations and predicted by the Flory–Huggins (FH) model. Labels indicate chain pairs. Simulated coefficients are normalized by (*R*_*i*_ + *R*_*j*_)^3^, where *R*_*i*_ is the radius of gyration of chain *i* in isolation, while predicted coefficients are normalized by the pervaded volume of an ideal polymer with chain length *L*_max_ [38]. (B) Correlation between the excess chemical potential differences measured in direct-coexistence simulations and predicted by the FH model. Points are shown for individual species in coexisting condensed phase pairs, *α*–*β*. The Pearson correlation coefficient, *ρ*, is shown for each panel.

By contrast, we find strong evidence that the pairwise-additive approximation holds to a greater extent in the condensed phases. We extract the excess chemical potential differences, 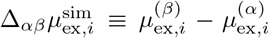, where the excess chemical potential is *µ*_ex,*i*_≡*µ*_*i*_−*µ*_id,*i*_, from the equilibrium compositions of simulated *α* and *β* bulk phases [25] and compare with the Flory–Huggins prediction, 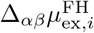(see SI). These measurements, which directly probe the accuracy of the pairwise approximation, Eq. (1), in the condensed phases, result in a much higher Pearson correlation coefficient of 0.93 (Fig. 4B). This result suggests that the polymer–polymer interactions are more mean-field-like in condensed phases, as originally pointed out by Flory [52]. More specifically, this observation suggests that the pairwise approximations for 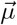, Eq. (1), and *ϵ* (Sec. II D) work well because the overlap parameter of our LJ heteropolymers is large in the condensed phases (*P* ≈ 13 *±* 2) [38].

**FIG. 5.**
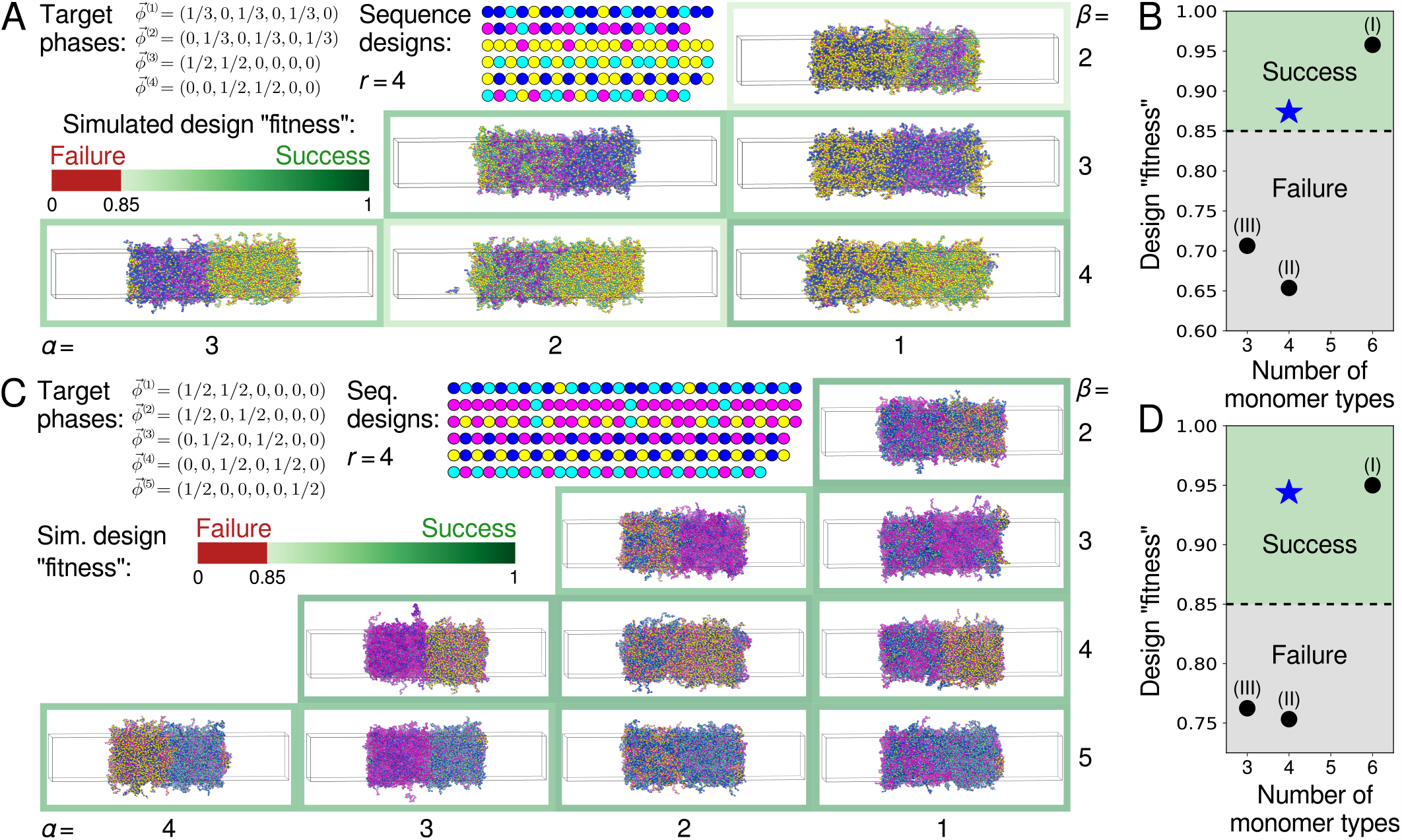
Design of complex multicomponent, multiphase condensates. (A) Inverse design and simulation validation of a 6-component, 4-condensed-phase system with the indicated target-phase molecular volume fractions. Sequence designs use the minimal required number of monomer types, *r* = 4. Equilibrium configurations of *α*–*β* direct-coexistence simulations are shown as in Fig. 3A, and the fitness metric is indicated by the border color. (B) Comparison of the optimized design shown in A with alternative designs, as in Fig. 3B. The alternative designs (I) maximize the stability of the condensed phases using six monomer types, (II) apply the EYM theorem without nuclear-norm minimization, and (III) apply NMF with *r* = 3 monomer types. (C) Inverse design and simulation validation of a 6-component, 5-condensed-phase system with the indicated target-phase molecular volume fractions. Here, the minimal required number of monomer types is also *r* = 4. (D) Comparison of the optimized design shown in C with alternative designs. Complete data for all designs are provided in the SI.

In line with our analysis of the excess chemical potentials, we find that the microstructures of the condensed phases agree with expectations based on the Gaussian Core Model (GCM) of finite-length polymers [53]. From simulations of individual, homogeneous condensed phases, we obtain the radial distribution function (RDF) for the center of mass of the polymer chains (see SI). Consistent with the GCM, the RDFs for all polymer species pairs exhibit a correlation hole, meaning that the centers of mass overlap with low probability, while a positive peak is found at a distance of approximately twice the radius of gyration, implying effective attractions between chains. The amplitudes of these peaks anticorrelate with the predicted pairwise interaction strengths given by *ϵ*. Most tellingly, the zero-wavenumber structure factor, which can be directly related to the pairwise interactions [37] (see SI), exhibits a Pearson correlation coefficient of 0.997 with respect to the predictions of the pairwise approximation. Thus, overall, our simulations provide a convincing *post hoc* validation of the key approximations utilized in applying our inverse design strategy to sequence-dependent LJ heteropolymers.

### G. Designing minimal-complexity biomolecular mixtures with complex phase behavior

The theory and inverse design approach that we describe here are broadly applicable for understanding complex, multiphase coexistence in biomolecular mixtures. To demonstrate its generality, we apply our inverse design approach to two additional interesting design problems, and then verify our predictions using simulations of LJ heteropolymers (see SI for complete details).

First, we consider a 6-component mixture in which components are shared among four condensed phases (Fig. 5A). This design problem is interesting because the number of enriched components differs among the condensed phases. Following the approach described in Sec. II D, we predict that at most *r* = 4 monomer types are required to construct LJ heteropolymers that phase separate into the prescribed phases. Performing simulations of all pairs of condensed phases as described in Sec. II E confirms this prediction, where we use the previously described fitness metric to assess the result of each phase pair. We also consider alternative designs in Fig. 5B, where we compare the minimum fitness of all phase pairs. These results are consistent with the predicted minimum number of monomer types and highlight the importance of following our inverse design algorithm to obtain a successful minimal-complexity design.

Second, we examine a scenario in which a 6-component mixture forms five condensed phases with shared components (Fig. 5C). Surprisingly, we find that the required number of molecular features, *r* = 4, is not only smaller than the number of components, but is also smaller than the number of condensed phases. To solve the matrix factorization problem, Eq. (6), we find that it is necessary to increase the maximum degree of polymerization to *L*_max_ = 30. Our simulations of designed LJ heteropolymers (Fig. 5C) and comparisons to alternative designs (Fig. 5D) similarly support our prediction that at most four monomer types are required to achieve the desired phase behavior. Taken together, these additional examples demonstrate the generality of our design approach and suggest that scenarios in which complex phase behavior can be realized using a small number of molecular features are not particularly rare.

## III. DISCUSSION

Our accomplishments in this paper are twofold. First, we establish a relationship between multiphase condensates and the complexity of the pairwise interactions in a biomolecular mixture. Specifically, we show that achieving multiphase organization via phase separation requires a minimal number of distinct molecular features, which represent linearly independent modes of interaction in a biomolecular mixture. While the physicochemical nature of these features may differ between different types of biomolecules, such as disordered proteins [54], nucleic acids [55], and multi-domain proteins [56], the required number of molecular features is determined solely by the target phase behavior. Second, we develop a two-step inverse design algorithm for calculating an upper bound on the required number of molecular features and then constructing minimal-complexity biomolecular mixtures that achieve the target phase behavior. We apply this inverse design approach to construct mixtures of model Lennard-Jones (LJ) heteropolymers that phase-separate into nontrivial coexisting phases, which would be extremely difficult to obtain without a rational design algorithm. Extensive molecular dynamics simulations provide direct support for the validity of our theory and inverse design approach, as well as justification for the additional approximations employed when applying our design approach to the LJ heteropolymer model.

Our results demonstrate that complex multiphase organization can be achieved using a relatively small number of chemical building blocks. On the one hand, this observation should not be surprising given the diversity of compositionally distinct phase-separated condensates that have been observed in living cells. Yet on the other hand, the way in which relatively “promiscuous” interactions among conformationally disordered biomolecules give rise to many immiscible condensates with controlled molecular compositions has been unclear until now. We speculate that the difference between the number of components in a mixture and the required number of molecular features will only tend to increase as the number of the components and the number of condensed phases grow. We reiterate that the number of monomer types computed in our studies of LJ heteropolymers represents an *upper bound* on the required number of monomer types, since the number of molecular features that govern interactions between linear polymers can exceed the number of monomer types. For example, it is well known that sequence-dependent properties, such as sequence “block-iness” [17, 57] and charge-patterning effects [58– tune the phase behavior of heteropolymer solutions in ways that cannot be predicted the monomer composition of the heteropolymers alone. In future work, we will incorporate such sequence-dependent features into our inverse design approach in order to tighten the bound on the required number of monomer types and broaden the range of applicability of our heteropolymer design algorithm. Overall, we hope that our approach will provide a unifying framework for recent efforts to identify molecular “grammars” governing phase separation in specific biomolecular systems [14–18].

Our two-step inverse design algorithm also establishes an extensible framework for designing phase-separating multicomponent fluids using a wide variety of synthetic building blocks. The key requirement is specifying the system-specific constraints on the molecular feature matrix and the feature–feature interactions. For example, we could apply our theory to design “patchy” colloidal particles [62], in which case the feature matrix indicates the surface area covered by a sticker type and the feature– feature interactions represent the interactions between different types of stickers. Nonetheless, we caution that making simplifying assumptions regarding the molecular interactions, such as the pairwise-additive approximation employed for LJ heteropolymers, are most likely to work well in systems with large overlap parameters; otherwise, the accuracy of the design algorithm is likely to be sensitive to the definitions of the molecular features and the way in which the molecular interactions are modeled.

In conclusion, our theory and inverse design approach advance our understanding of multiphase condensate formation in biology, as well as our ability to engineer chemically diverse artificial condensates. Our predictions could be tested using existing experimental technologies in both biological [54, 63–67] and synthetic [68–71] systems. Ultimately, by incorporating the design of tunable interfacial tensions [72, 73] and nonequilibrium chemical activity [74, 75] into our framework, it will be possible to realize fully “programmable” fluids and soft materials that self-organize via multicomponent phase separation.

## ACKNOWLEDGMENTS

The authors acknowledge Ofer Kimchi, Isabella Graf, and Zhuang Liu for providing comments on a previous version of this manuscript and Princeton Research Computing for technical support. Source code and example calculations can be found at https://github.com/wmjac/phaseprogramming-pub.

## Notes

### Competing Interest Statement

The authors have declared no competing interest.

