## Supplementary Information for "Emergence of multiphase condensates from a limited set of chemical building blocks"

### I. RELATING INTERACTION INTERDEPENDENCIES AND BIOMOLECULAR CONDENSATE THERMODYNAMICS

#### A. Multicomponent macromolecular solutions with pairwise interactions

Throughout this work, we consider a solution of $N$ macromolecular species in an implicit solvent. The concentrations (i.e., number densities) of the macromolecular species, $\{\rho_i\}$, are related to the macromolecular volume fractions, $\{\phi_i \equiv v_i\rho_i\}$, by the macromolecular excluded volumes, $\{v_i\}$. As discussed in the main text, we assume a free-energy density $f$ in which the molecular interactions follow a pairwise approximation. This means that the (scaled) chemical potentials of all macromolecular components, $\mu_i = \partial f/\partial\phi_i = v_i^{-1}\partial f/\partial\rho_i$, can described by the equation

$$\mu_i(\vec{\phi}) = \mu_{\mathrm{id},i}(\vec{\phi}) + \mu_{\mathrm{v},i}(\vec{\phi}) + \sum_{j=1}^{N} \epsilon_{ij}\phi_j, \tag{S1}$$

where $\mu_{\mathrm{id},i} = v_i^{-1}k_{\mathrm{B}}T\log\phi_i$ is the ideal contribution to the chemical potential and $\mu_{\mathrm{v},i}(\vec{\phi})$ accounts for steric contributions. Note that the chemical potential of macromolecular species $i$, as defined above, is scaled with respect to the excluded volume $v_i$; this choice is made for convenience and does not affect our results. Both the ideal and the steric contributions are independent of the pairwise interaction matrix, $\boldsymbol{\epsilon}$. Meanwhile, the pairwise interaction matrix, $\boldsymbol{\epsilon}$, is assumed to be independent of the macromolecular concentrations.

#### B. Perturbative analysis of the landscape stability

Applying small perturbations to the component-wise interactions $\{\epsilon_{ij}\}$, we expand the free-energy density, $f$, to linear order,

$$\tilde{f} = f + \sum_{i=1}^{N}\sum_{j=i}^{N} \frac{\partial f}{\partial\epsilon_{ij}}\Delta\epsilon_{ij} + \sum_{i=1}^{N}\frac{\partial f}{\partial\phi_i}\Delta\phi_i. \tag{S2}$$

Since $\boldsymbol{\epsilon}$ is a symmetric $N\times N$ matrix, we only consider the independent degrees of freedom $\{\epsilon_{ij}\}$ for which $i \le j$. Eq. (S1) implies that $\partial f/\partial\epsilon_{ij} = (1/2)^{\delta_{ij}}\phi_i\phi_j$. We can therefore rewrite the term in Eq. (S2) involving $\Delta\epsilon_{ij}$ in matrix–vector notation,

$$\begin{aligned}\sum_{i=1}^{N}\sum_{j=i}^{N}\frac{\partial f}{\partial\epsilon_{ij}}\Delta\epsilon_{ij} &= \frac{1}{2}\Bigg(\sum_{i=1}^{N}\phi_i\Delta\epsilon_{ii}\phi_i + 2\sum_{i<j}\phi_i\Delta\epsilon_{ij}\phi_j\Bigg)\\ &= \frac{1}{2}\Bigg(\sum_{i=1}^{N}\phi_i\Delta\epsilon_{ii}\phi_i + \sum_{i<j}\phi_i\Delta\epsilon_{ij}\phi_j + \sum_{i>j}\phi_i\Delta\epsilon_{ij}\phi_j\Bigg)\\ &= \frac{1}{2}\vec{\phi}^{\top}\Delta\boldsymbol{\epsilon}\vec{\phi}.\end{aligned} \tag{S3}$$

We then expand the chemical potentials at phase coexistence,

$$\tilde{\mu}_k = \mu_k + \sum_{j=i}^{N}\sum_{i=1}^{N}\frac{\partial^2 f}{\partial\epsilon_{ij}\partial\phi_k}\Delta\epsilon_{ij} + \sum_{i=1}^{N}\frac{\partial^2 f}{\partial\phi_k\partial\phi_i}\Delta\phi_i. \tag{S4}$$

The change in the chemical potentials, $\Delta\vec{\mu}$, can be written in matrix–vector notation as

$$\Delta\vec{\mu} \equiv \tilde{\vec{\mu}} - \vec{\mu} = \Delta\boldsymbol{\epsilon}\vec{\phi} + \left(\frac{\partial^2 f}{\partial\vec{\phi}^2}\right)\Delta\vec{\phi}. \tag{S5}$$

The grand potential density in the $\alpha$ phase, $\tilde{\Omega}^{(\alpha)} \equiv \tilde{f} - \tilde{\vec{\mu}} \cdot \vec{\phi}$, can similarly be written as

$$\begin{aligned}\tilde{\Omega}^{(\alpha)} &= \Omega^{(\alpha)} + \frac{\partial f}{\partial \vec{\phi}}\bigg|^{(\alpha)} \cdot \Delta\vec{\phi}^{(\alpha)} + \sum_{i=1}^{N}\sum_{j=i}^{N} \frac{\partial f}{\partial \epsilon_{ij}}\bigg|^{(\alpha)} \Delta\epsilon_{ij} - \vec{\mu} \cdot \Delta\vec{\phi}^{(\alpha)} - \Delta\vec{\mu}^{(\alpha)} \cdot \vec{\phi}^{(\alpha)} \\ &= \Omega^{(\alpha)} + \vec{\mu} \cdot \Delta\vec{\phi}^{(\alpha)} + \frac{1}{2}\vec{\phi}^{(\alpha)\top} \Delta\boldsymbol{\epsilon}\vec{\phi}^{(\alpha)} - \vec{\mu} \cdot \Delta\vec{\phi}^{(\alpha)} - \Delta\vec{\mu}^{(\alpha)} \cdot \vec{\phi}^{(\alpha)} \\ &= \Omega^{(\alpha)} + \frac{1}{2}\vec{\phi}^{(\alpha)\top} \Delta\boldsymbol{\epsilon}\vec{\phi}^{(\alpha)} - \Delta\vec{\mu}^{(\alpha)} \cdot \vec{\phi}^{(\alpha)}.\end{aligned} \tag{S6}$$

At coexistence, it is required that $\vec{\mu}^{(\alpha)} = \vec{\mu}^{(0)}$ and $\Omega^{(\alpha)} = \Omega^{(0)}$ for every condensed phase $\alpha = 1, \ldots, K$, where the index 0 indicates the dilute phase. Requiring that the perturbed phases remain at coexistence means that $\Delta\vec{\mu}^{(\alpha)} = \Delta\vec{\mu}^{(0)}$ and $\tilde{\Omega}^{(\alpha)} = \tilde{\Omega}^{(0)}$. Since $\Delta\vec{\mu} = \Delta\boldsymbol{\epsilon}\vec{\phi} + (\partial^2 f / \partial \vec{\phi}^2)\Delta\vec{\phi}$ from Eq. (S5), we find that the coexistence condition for $\tilde{\Omega}$ is

$$-\frac{1}{2}(\vec{\phi}^{(\alpha)} - \vec{\phi}^{(0)})^\top \Delta\boldsymbol{\epsilon}(\vec{\phi}^{(\alpha)} + \vec{\phi}^{(0)}) = (\vec{\phi}^{(\alpha)} - \vec{\phi}^{(0)})^\top \left(\frac{\partial^2 f}{\partial \vec{\phi}^2}\right)\Delta\vec{\phi}. \tag{S7}$$

Assuming that the dilute phase is very dilute, this result simplifies to

$$\Delta\boldsymbol{\epsilon}\vec{\phi}^{(\alpha)} = -2\frac{\partial^2 f}{\partial \vec{\phi}^2}\bigg|^{(\alpha)} \cdot \Delta\vec{\phi}. \tag{S8}$$

Eq. (S8) relates changes in the macromolecular concentrations in the $\alpha$ phase to perturbations in the pairwise interactions, assuming that the Hessian matrix of the $\alpha$ phase is known. We can then estimate the noise tolerance of the interaction matrix $\boldsymbol{\epsilon}$ in the worst-case scenario by considering compositional changes along the least stable direction, $\hat{\nu}_1^{(\alpha)}$, of each target phase, where

$$\frac{\partial^2 f}{\partial \vec{\phi}^2}\bigg|^{(\alpha)} \cdot \hat{\nu}_1^{(\alpha)} = \lambda_1^{(\alpha)}\hat{\nu}_1^{(\alpha)}, \tag{S9}$$

and the eigenvalues of the Hessian matrix are $0 > \lambda_1^{(\alpha)} \geq \lambda_2^{(\alpha)} \geq \ldots \geq \lambda_N^{(\alpha)}$. Projecting an arbitrary concentration change $\Delta\vec{\phi}$ onto the least stable direction, we find

$$\|\Delta\boldsymbol{\epsilon}\vec{\phi}^{(\alpha)}\|_F = \eta\lambda_1^{(\alpha)}, \tag{S10}$$

where $\eta \equiv 2\Delta\vec{\phi} \cdot \hat{\nu}_1^{(\alpha)} / \|\vec{\phi}^{(\alpha)}\|$ corresponds to twice the relative percentage change of macromolecular composition in the $\alpha$ phase and $\|\cdot\|_F$ is the Frobenius norm. Applying the sub-multiplicative property of the Frobenius norm to the left-hand side, we have

$$\|\Delta\boldsymbol{\epsilon}\hat{\phi}\|_F \leq \|\Delta\boldsymbol{\epsilon}\|_F \|\hat{\phi}\|_2;$$

the equality holds if $\hat{\phi}$ and each row of $\Delta\boldsymbol{\epsilon}$ are linearly independent. Assuming that this is the case, we can express the tolerance of the reconstruction error for each target phase $\alpha$ in terms of an allowed relative composition change $\eta$,

$$\|\Delta\boldsymbol{\epsilon}\|_F \approx \eta\lambda_1^{(\alpha)}. \tag{S11}$$

The overall tolerance is therefore given by

$$\|\Delta\boldsymbol{\epsilon}\|_F \approx \min_\alpha \eta\lambda_1^{(\alpha)}. \tag{S12}$$

Based on this analysis, the optimal interaction matrix $\boldsymbol{\epsilon}$ should simultaneously maximize the landscape stability and minimize the reconstruction error, $\|\Delta\boldsymbol{\epsilon}\|_F$. Using Eq. (S12), we can apply the Eckart–Young–Mirsky (EYM) theorem [1] to find that the smallest number of distinct molecular features, $r$, that successfully solve the inverse design problem,

$$\left[\sum_{k=1}^{N-r} \sigma_k^2\right]^{1/2} \lesssim \min_\alpha \eta\lambda_1^{(\alpha)}, \tag{S13}$$

where $\sigma_1 \leq \sigma_2 \leq \ldots \leq \sigma_N$ are the singular values of $\boldsymbol{\epsilon}$. In other words, for a given $N \times N$ interaction matrix $\boldsymbol{\epsilon}$, the rank-$r$ approximation, $\boldsymbol{\epsilon_r}$, that minimizes $\|\boldsymbol{\Delta\epsilon}\|_F = \|\boldsymbol{\epsilon_r} - \boldsymbol{\epsilon}\|_F$ can be obtained by eliminating the smallest $N - r$ singular values of $\boldsymbol{\epsilon}$. If these smallest singular values are nonzero, then the left-hand-side of Eq. (S13) will be nonzero, and the rank-$r$ approximation of the interaction matrix, $\boldsymbol{\epsilon_r}$, will not exactly equal the original interaction matrix, $\boldsymbol{\epsilon}$. However, the macromolecular compositions of the resulting coexisting phases will deviate from the target compositions within the allowed tolerance $\eta$ if Eq. (S13) is satisfied. Eq. (S13) thus motivates the identification of the minimal number of distinct molecular features, $r$, needed to stabilize a set of target phases.

### II. TWO-STEP OPTIMIZATION-BASED DESIGN APPROACH

#### A. Convex relaxation for component-wise interactions

In order to obtain coexisting phases with prescribed macromolecular volume fractions $\{\vec{\phi}^{(1)}, \vec{\phi}^{(2)}, \ldots, \vec{\phi}^{(K)}\}$, we first solve for the component-wise interactions using convex programming. We refer the reader to Ref. [2] for a detailed discussion of this approach. The convex relaxation of the thermodynamic-stability and target-volume-fraction constraints defines a semi-definite program (SDP) [3],

$$\mu_{\mathrm{id},i}(\vec{\phi}^{(\alpha)}; \vec{v}) + \mu_{\mathrm{ex},i}(\vec{\phi}^{(\alpha)}; \boldsymbol{\epsilon}, \vec{v}) \geq \mu_i \;\forall i, \alpha \tag{S14a}$$

$$P(\vec{\phi}^{(\alpha)}; \boldsymbol{\epsilon}, \vec{v}) = 0 \;\forall \alpha \tag{S14b}$$

$$\partial[\vec{\mu}_{\mathrm{id}}(\vec{\phi}^{(\alpha)}; \vec{v}) + \vec{\mu}_{\mathrm{ex}}(\vec{\phi}^{(\alpha)}; \boldsymbol{\epsilon}, \vec{v})]/\partial\vec{\phi} \succ \lambda_{\min} I \;\forall \alpha \tag{S14c}$$

$$\phi_{\mathrm{T}}^{(0)}(\vec{\mu}; \vec{v}) < \phi_{\mathrm{T}}^{*}(\vec{v}). \tag{S14d}$$

Components are either "enriched" or "depleted" in the $\alpha$ phase, depending on whether they make a substantial or negligible contribution, respectively, to the macromolecular composition of that phase. The ideal chemical potential is $\mu_{\mathrm{id},i} = v_i^{-1} \log \phi_i^{(\alpha)}$ for any component $i$ that is enriched in the $\alpha$ phase or $\mu_{\mathrm{id},i} = v_i^{-1} \log \phi_{\mathrm{depl}}^{(\alpha)}$ for any component $i$ that is depleted in the $\alpha$ phase. The equality(inequality) in Eq. (S14a) applies to enriched(depleted) components. The volume fraction of any component that is depleted in the $\alpha$ phase cannot exceed $\phi_{\mathrm{depl}}^{(\alpha)} \equiv \zeta \phi_{\mathrm{T}}^{(\alpha)}/[M^{(\alpha)}(N - M^{(\alpha)})]$, where $\phi_{\mathrm{T}}^{(\alpha)}$ is the total volume compositions of all components in the $\alpha$ phase, $M^{(\alpha)}$ is the number of enriched components in the $\alpha$ phase, and $\zeta$ is an adjustable parameter. In this work, we choose $\phi_{\mathrm{T}} = 0.9$ and $\zeta = 10^{-2}$ for all target phases. Eq. (S14b) states that the pressure, $P$, must be zero in all phases, since the pressure in the dilute phase, which is nearly ideal, is also approximately zero. In Eq. (S14c), the parameter $\lambda_{\min} \geq 0$ places a lower bound on the smallest eigenvalue of the Hessian matrix in order to guarantee thermodynamic stability. The final constraint, Eq. (S14d), ensures that the volume fraction in the dilute phase, $\phi_{\mathrm{T}}^{(0)}$, is less than the critical volume fraction, $\phi_{\mathrm{T}}^{*}(\vec{v})$. This condition is independent of $\boldsymbol{\epsilon}$ due to the zero-osmotic-pressure assumption. We implement and solve this SDP in practice using efficient convex-optimization software [4, 5].

Within the approximations of this convex relaxation, the constraints given by Eq. (S14) define the joint space of interaction matrices, $\boldsymbol{\epsilon}$, and chemical potential vectors, $\vec{\mu}$, for which bulk phase coexistence can be established among the target condensed phases and a dilute phase. Based on the perturbative analysis in SI Sec. I B, we choose to minimize the nuclear norm, $\|\boldsymbol{\epsilon}\|_* \equiv \sum_{i=1}^{N} \sigma_i$, to pick out a unique solution from within the solution space that satisfies the constraints, Eq. (S14). This objective function is a convex relaxation of the matrix rank [6] and thus tends to select $\boldsymbol{\epsilon}$ matrices with small singular values. We then identify the smallest number of molecular features, $r$, with which we can satisfy the bound given in Eq. (S13). Finally, we obtain the low-rank approximation $\boldsymbol{\epsilon_r}$ for the component-wise interactions via the EYM theorem. By contrast, the objective function used in Ref. [2] tends to select $\boldsymbol{\epsilon}$ with larger singular values; as a result, solutions obtained using this alternative objective function tend to require a larger number of molecular features to satisfy the bound given in Eq. (S13).

#### B. Stochastic optimization for feature-wise interactions

In the second step of our design approach, we aim to factorize the low-rank approximation of the component-wise interaction matrix, $\boldsymbol{\epsilon_r}$, by minimizing the reconstruction error $\|\Delta^{\mathrm{recon}}\epsilon\|_F$. However, we are, in general, unable to use a straightforward eigenvalue decomposition of $\boldsymbol{\epsilon_r}$, since we typically wish to interpret the factorization in terms of nonnegative molecular feature vectors. This means that we must, in general, solve a nonnegative matrix factorization (NMF) problem [7]. To this end, we specify the loss function

$$\mathcal{L} \equiv \|\Delta^{\mathrm{recon}}\epsilon\|_F^2 = \|\boldsymbol{\epsilon_r} - \boldsymbol{W}\boldsymbol{u}\boldsymbol{W}^\top\|_F^2, \tag{S15}$$

which is nonlinear with respect to the independent variables $\boldsymbol{W}$ and $\boldsymbol{u}$. Here we consider two possible scenarios, in which the molecular feature matrix, $\boldsymbol{W}$, can either take nonnegative real values or nonnegative integers.

##### 1. *NMF without integer constraints*

When there are no integer constraints on the molecular feature matrix $\boldsymbol{W}$, Eq. (S15) is biconvex with respect to either $\boldsymbol{W}$ or $\boldsymbol{u}$. The NMF problem can therefore be solved by iteratively alternating optimization of $\boldsymbol{W}$ and $\boldsymbol{u}$, with guaranteed convergence to a local optimum [7]. Here we consider a scenario in which $\boldsymbol{W}$ takes positive real values (representing, e.g., the fraction of the surface area of a colloidal particle covered by "patches" of certain types [8]) and $\boldsymbol{u}$ takes negative real values (representing, e.g., the attractive interactions between patches). To ensure nonnegativity of $\boldsymbol{W}$ and nonpositivity of $\boldsymbol{u}$, we apply a multiplicative update scheme, where $\boldsymbol{W}$ and $-\boldsymbol{u}$ are multiplied by the ratio of the negative and positive parts of the corresponding gradient, $\nabla = \nabla^+ - \nabla^-$. We then let

$$W_{ia} \leftarrow W_{ia}\left(\frac{\nabla W_{ia}^-}{\nabla W_{ia}^+}\right), \tag{S16}$$

$$-u_{ab} \leftarrow -u_{ab}\left(\frac{\nabla(-u)_{ab}^-}{\nabla(-u)_{ab}^+}\right). \tag{S17}$$

This scheme is equivalent to gradient descent with a step size proportional to the ratio $\nabla^-/\nabla^+$. In the absence of any additional constraints, the update rules are thus

$$W_{ia} \leftarrow W_{ia}\left(\frac{(\boldsymbol{\epsilon}\boldsymbol{W}\boldsymbol{u}^\top + \boldsymbol{\epsilon}^\top\boldsymbol{W}\boldsymbol{u})_{ia}}{(\boldsymbol{W}\boldsymbol{u}\boldsymbol{W}^T\boldsymbol{W}\boldsymbol{u}^\top + \boldsymbol{W}\boldsymbol{u}^\top\boldsymbol{W}^\top\boldsymbol{W}\boldsymbol{u})_{ia}}\right)^\gamma, \tag{S18}$$

$$-u_{ab} \leftarrow -u_{ab}\left(\frac{(\boldsymbol{W}^\top\boldsymbol{\epsilon}\boldsymbol{W})_{ab}}{(\boldsymbol{W}^\top\boldsymbol{W}\boldsymbol{u}\boldsymbol{W}^\top\boldsymbol{W})_{ab}}\right)^\gamma, \tag{S19}$$

where we choose $\gamma = 1/4$ to ensure stable convergence. We can also include additional constraints, such as minimizing the variance of all monomer–monomer interactions or the variance of homotypic monomer–monomer interactions. These constraints can be easily implemented by adding the corresponding positive and negative parts of the constraints to the gradient, and then following the update scheme given above.

#### 2. *NMF with integer constraints*

Imposing integer constraints on $\boldsymbol{W}$ makes the NMF problem no longer convex in $\boldsymbol{W}$. Thus, in cases such as the heteropolymer design problem considered in the main text, where the molecular feature matrix must have integer entries (see also SI Sec. III A), optimizing $\boldsymbol{W}$ becomes an NP-hard combinatorial optimization problem. For short polymer chains with $L_{\max} \lesssim 10$ and a prescribed monomer–monomer interaction matrix $\boldsymbol{u}$, it is feasible to find the globally optimal $\boldsymbol{W}$ matrix by brute force. However, brute-force search is infeasible for designing long polymer chains. Instead, we follow an approach in which we sample $\boldsymbol{W}$ matrices from a probability distribution, and then select candidate $\boldsymbol{W}$ matrices from the tail of the reconstruction-error distribution. Suppose that every row of the $\boldsymbol{W}$ matrix is drawn from a multinomial distribution $\vec{W}_i \sim \mathcal{M}_r(L_{\max}; \vec{p}_i)$ for $i = 1, \ldots, N$, which gives the probability of any particular combination of counts for various feature types $a = 1, \ldots, r$. The probability vector $\vec{p}_i$, which is normalized such that $\sum_{a=1}^r p_{ia} = 1$, parameterizes the multinomial distribution. In the heteropolymer design scenario, the number of categorical variables $r = \operatorname{rank}(\boldsymbol{\epsilon}_r)$ denotes the number of monomer types, $L_{\max}$ is equal to the sum of the counts in $\vec{W}_i$, and $p_{ia}$ represents the probability of adding a monomer of type $a$ to chain $i$.

We are interested in finding an integer $\boldsymbol{W}$ solution in which the reconstruction error is smaller than a prescribed threshold. If such $\boldsymbol{W}$ matrices are rare, then we can carry out the search using a cross-entropy (CE) optimization approach [9]. This algorithm proceeds as follows:

1. Choose an initial parameter matrix $\boldsymbol{p}$, which contains all the probability vectors $\vec{p}_i$. Choose the total number of samples to be generated ($N_{\text{samples}} = 10000$) and the number of samples needed for parameter inference ($N_{\text{top}} = 10$). Let the initial iteration number be $t = 1$.

2. To generate a candidate solution pair $(\boldsymbol{W}, \boldsymbol{u})$, sample each row $\vec{W}_1, ..., \vec{W}_N \sim_{\text{iid}} \mathcal{M}_r(L_{\max}, \vec{p}_i)$. Then, given this candidate $\boldsymbol{W}$ matrix, use gradient descent to optimize for a nonpositive $\boldsymbol{u}$ matrix by iteratively applying the multiplicative update given in Eq. (S19) until convergence, defined as the point where the maximum difference between updates of any element of $\boldsymbol{u}$ is less than $10^{-5}$.

3. Repeat step 2 to generate $N_{\text{samples}}$ samples and calculate the loss, Eq. (S15), for each sample.

4. Update $\boldsymbol{p}^{t+1} \leftarrow \boldsymbol{p}^t$ according to a maximum likelihood estimate. Specifically, determine the probability vectors $\vec{p}_i^{\,t+1}$ by computing the normalized average frequencies of all monomer types observed in chain type $i$ in the $N_{\text{top}}$ candidate solutions with the smallest values of the loss.

5. Iterate steps 2, 3, and 4 until the maximum difference between updates of any element of the parameter matrix $\boldsymbol{p}$ is less than 0.005 or the maximal number of iterations (chosen here to be 100) is exceeded. The $(\boldsymbol{W}, \boldsymbol{u})$ pair with the minimal loss is considered to be a solution if the reconstruction error is below the threshold, Eq. (S12).

We use uniform random initialization of the probability vectors to obtain all the design solutions presented in this paper. Because permutation of the rows of $\boldsymbol{W}$ could potentially lead to degenerate solutions during the search, we sort the rows with respect to descending monomer frequencies for every candidate $\boldsymbol{W}$ matrix. Furthermore, to make the optimization problem easier, we relax the constraint on the degree of polymerization by allowing the sequence length to be shorter than $L_{\max}$. To this end, we allow $\sum_{a=1}^{r} p_{ia} \leq 1$, which is realized in practice by adding a dummy column to the probability matrix $\boldsymbol{p}$. This enhanced-sampling method allows us to efficiently search for NMF solutions for arbitrary long polymer chains.

### III. MULTICOMPONENT POLYMER MODEL AND SIMULATION METHODS

#### A. Multicomponent Flory–Huggins polymer model

We utilize the mean-field Flory–Huggins (FH) model to apply our theory to the design of Lennard–Jones heteropolymer solutions. The Helmholtz free-energy density, $f$; chemical potential, $\vec{\mu} = \partial f / \partial \vec{\phi}$; osmotic pressure, $P$; and Hessian matrix, $\partial^2 f / \partial \vec{\phi}^2 = \partial \vec{\mu}(\vec{\phi}) / \partial \vec{\phi}$, in the multicomponent Flory–Huggins model are

$$f = \sum_{i=1}^{N} \frac{\phi_i}{L_i} \log \phi_i + (1 - \phi_{\mathrm{T}}) \log(1 - \phi_{\mathrm{T}}) + \frac{1}{2} \sum_{i=1}^{N} \sum_{j=1}^{N} \epsilon_{ij} \phi_i \phi_j \tag{S20a}$$

$$\mu_i = \frac{1}{L_i} \log \phi_i - \log(1 - \phi_{\mathrm{T}}) - \left(1 - \frac{1}{L_i}\right) + \sum_{j=1}^{N} \epsilon_{ij} \phi_j \tag{S20b}$$

$$P = -\log(1 - \phi_{\mathrm{T}}) + \sum_{i=1}^{N} \frac{\phi_i}{L_i} - \phi_{\mathrm{T}} + \frac{1}{2} \sum_{i=1}^{N} \sum_{j=1}^{N} \epsilon_{ij} \phi_i \phi_j \tag{S20c}$$

$$\frac{\partial \mu_i}{\partial \phi_j} = \frac{\delta_{ij}}{L_i \phi_i} + \frac{1}{1 - \phi_{\mathrm{T}}} + \epsilon_{ij}, \tag{S20d}$$

respectively, where $L_i$ is the degree of polymerization of polymeric species $i$. When applying our theory and design approach to multicomponent polymer solutions, we use these expressions in the convex relaxation, Eq. (S14).

When designing heteropolymers within this mean-field framework, we make a pairwise-additive assumption for the polymer–polymer interactions, $\boldsymbol{\epsilon}$. This means that the interaction matrix can be written as $\epsilon_{ij} = \sum_{a,b} W_{ia} u_{ab}^{\mathrm{LJ}} W_{jb}$, where the molecular feature matrix, $\boldsymbol{W}$, is a count encoding of the occurrences of each monomer type, $a = 1, \ldots, r$, within each polymer chain type, $i = 1, \ldots, N$. This factorization of $\boldsymbol{\epsilon}$ follows directly from the Flory–Huggins model, since the pairwise portion of the free-energy density, $(1/2) \sum_{i,j} \epsilon_{ij} \phi_i \phi_j$, represents the mean-field interaction between all pairs of monomers in a polymer solution [10]. More specifically, we can write this sum in the factorized form $(1/2) \sum_{i,j} \sum_{a,b} u_{ab} W_{ia} W_{jb} \phi_i \phi_j = (1/2) \sum_{a,b} u_{ab} \tilde{\phi}_a \tilde{\phi}_b$, where $\tilde{\phi}_a \equiv \sum_i W_{ia} \phi_i$ is the volume fraction occupied by monomers of type $a$. As we also emphasize in the main text, this pairwise-additive approximation for the polymer–polymer interactions is not an essential assumption of our theory, since we could directly determine the molecular features of heteropolymer interactions from simulations and then use these inferred features as a basis set for designing heteropolymer mixtures with prescribed polymer–polymer interactions. Such an approach could potentially capture sequence-patterning effects, which are explicitly ignored by the pairwise-additive assumption. (This topic will be the subject of future work.) Here we employ the pairwise-additive assumption because the meaning of the $\boldsymbol{W}$ matrix is easy to understand in this case, thus providing a transparent proof-of-principle for our theory. We also show (see SI Sec. III E) that this approximation is surprisingly accurate for LJ heteropolymers under our simulation conditions.

#### B. Lennard–Jones heteropolymer model

##### 1. *Simulation model*

We perform molecular simulations of heteropolymers in implicit solvent using a Lennard–Jones (LJ) polymer model using the LAMMPS simulation package [11]. Nonbonded monomers of types $a, b = 1, \ldots, r$ interact via the LJ pair potential [12],

$$U_{ab}^{\mathrm{LJ}}(r) = 4w_{ab}^{\mathrm{LJ}}\left[(d/r)^{12} - (d/r)^{6}\right] + U_{\mathrm{cut},ab}, \tag{S21}$$

where $w_{ab}^{\mathrm{LJ}} < 0$ represents the interaction strength (i.e., well depth), $d$ is the monomer diameter, and the cutoff distance is $3d$. The potential is shifted to zero at the cutoff distance by setting $U_{\mathrm{cut},ab} = -4w_{ab}^{\mathrm{LJ}}[(1/3)^{12} - (1/3)^{6}]$. Bonded monomers interact via finite-extensible nonlinear elastic (FENE) bonds [13], which comprise a nonlinear attractive term and a repulsive LJ term. For the FENE bonds, we use the parameters $(100, 4, 1, 1)$, corresponding to the coefficient of the attractive term, the maximum length of the bond in units of $d$, the well depth for the LJ potential, and the diameter of the particle in units of $d$.

##### 2. *Constructing LJ interaction parameters and heteropolymer sequences*

To perform simulation tests of our heteropolymer mixture designs, we need to construct LJ interaction matrices, $\{w_{ab}^{\mathrm{LJ}}\}$, and polymer sequences using the optimal $\boldsymbol{u}^{\mathrm{LJ}}$ and $\boldsymbol{W}$ matrices that we obtain via NMF (see Sec. II B 2). We describe each of these steps in this section.

First, to convert between the mean-field monomer–monomer interaction coefficients, $\{u_{ab}^{\mathrm{LJ}}\}$, and the well depths for the LJ interactions, $\{w_{ab}^{\mathrm{LJ}}\}$, we perform a nonlinear mapping by matching the monomer–monomer second virial coefficients. Specifically, we approximate the repulsive part of the LJ potential using the hard-sphere potential, and then relate the attractive portion of the LJ second virial coefficient to $u_{ab}^{\mathrm{LJ}}$,

$$\frac{u_{ab}^{\mathrm{LJ}}}{|\bar{u}^{\mathrm{LJ}}|} = \frac{2\pi}{d^3}\int_d^{3d} dr\, r^2 \left\{1 - \exp[-U_{ab}^{\mathrm{LJ}}(r)/k_{\mathrm{B}}T]\right\}, \tag{S22}$$

given a fixed temperature $T$. In practice, we work at a standard temperature, such that $k_{\mathrm{B}}T = 1$ in our simulations. The user-defined scale factor, $|\bar{u}^{\mathrm{LJ}}|$, controls the mean LJ interaction strengths used in the simulations; this scale factor must be introduced empirically (as opposed to being determined from the mean-field model) because the critical points of the mean-field FH model and the LJ heteropolymer simulations differ. In practice, we determine the LJ coefficients $\{w_{ab}^{\mathrm{LJ}}\}$ from a designed monomer–monomer interaction matrix, $\boldsymbol{u}^{\mathrm{LJ}}$, by choosing an appropriate value of $|\bar{u}^{\mathrm{LJ}}|$ (determined from simulations of homomeric LJ heteropolymers with chain length $L_{\max}$) and inverting Eq. (S22).

Second, to design the polymer sequences for polymer species $i = 1, \ldots, N$, we utilize the count encodings in each row, $\vec{W}_i$, of the optimized molecular feature matrix. We aim to construct sequences that are minimally "blocky", since the pairwise-additive assumption used in our heteropolymer design approach (see Sec. III A) does not consider sequence-patterning effects. We therefore aim to construct sequences in which the monomers of each type are homogeneously distributed throughout each polymer chain. Here we use a deterministic heuristic for interleaving different monomer types. To generate the sequence design for chain $i$, we first sort the monomer frequency counts $\{W_{ia}\}$ in descending order and associate the $n$th monomer of type $a$ with the fractional number $n/(1 + W_{ia})$, where $n$ goes from 1 to $W_{ia}$. We then read off the order of the monomer types in the sequence by going through the array of fractional numbers in ascending order.

Following these procedures, we obtain the LJ coefficients used for simulating the heteropolymer design shown in Fig. 3C in the main text,

$$\boldsymbol{w}^{\mathrm{LJ}} = \begin{bmatrix} 0.520 & 0.863 & 0.380 \\ 0.863 & 0.345 & 0.238 \\ 0.380 & 0.238 & 0.788 \end{bmatrix}.$$

The corresponding sequences are shown in Fig. 3C of the main text.

#### C. Direct-coexistence simulations

##### 1. *Initialization of multiphase simulations*

To prepare condensed phases for molecular dynamics simulations, we first perform constant-temperature-and-pressure (NPT) simulations for each individual target phase. We use $N_{\mathrm{tot}} = 432$ chains in a cubic box with periodic boundary conditions, where chain types are assigned according to the target-phase composition. Performing NPT

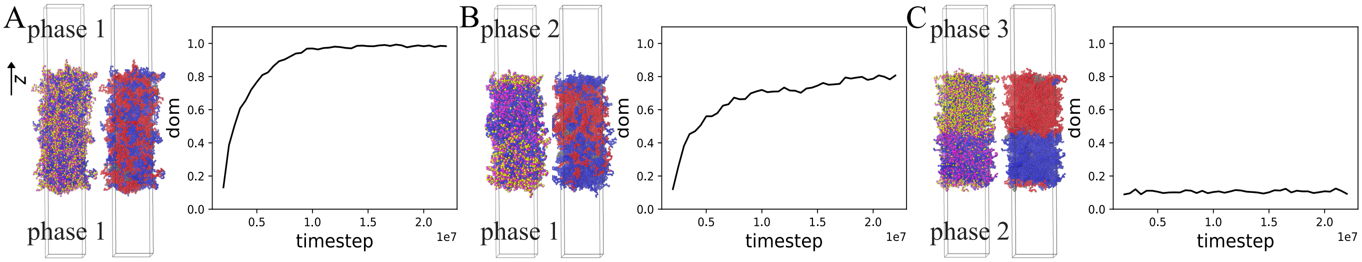


FIG. S1. **The degree of mixing as an indicator of equilibration.** For the example design solution shown in Fig. 3 in the main text, we compare the evolution of the degree of mixing (d.o.m.) between (A) an $\alpha = \beta$ control simulation and (B–C) two $\alpha \neq \beta$ direct-coexistence simulations. The left and right snapshots in each panel are colored according to the monomer types and the initial phase labels (either red or blue) at the end of the simulation trajectory. From **A**, we determine that the mixing timescale at this temperature and mean LJ interaction strength is approximately $10^7$ timesteps ($5 \times 10^4 \tau$). The d.o.m. is significantly greater than zero in **B** because phases 1 and 2 share an enriched component. Interfacial effects, both at the $\alpha$–$\beta$ interface and at the condensed–dilute interfaces, also tend to increase the d.o.m. in both **B** and **C**.

simulations at zero pressure allows us to estimate the average equilibrium total number density in a condensed phase, $\rho_{\text{eq}}$. We then prepare the condensed phases for use in direct-coexistence simulations by deforming the simulation box. In this step, we maintain periodic boundary conditions in both the x and y directions while imposing hard wall constraints and open boundary conditions in the z direction. The final dimensions, consistent with the target density $\rho_{\text{eq}}$ determined from the NPT simulations, are $l_z = 40d$ and $l_x = l_y = 20d$ for simulations of designs with $L_{\max} = 20$, and $l_z = 40d$ and $l_x = l_y = 24.5d$ for simulations of designs with $L_{\max} = 30$. Finally, the initial condition for a direct-coexistence simulation of a pair of condensed phases is constructed by stitching together two condensed phases and a dilute gas phase along the z axis of the simulation box. The region of the simulation box corresponding to the dilute phase is initialized with 6 chains of each type at a total number density of $\rho_{\text{dilute}} = 1.50 \times 10^{-3}$, $2.25 \times 10^{-3}$, or $1.50 \times 10^{-3}$ for the designs shown in Fig. 3A, Fig. 5A, and Fig. 5C, respectively.

#### *2. Production simulations*

In our production runs, we perform constant-temperature-and-volume (NVT) direct-coexistence simulations. The overall dimensions of the simulation box, including $\alpha$, $\beta$, and dilute phases, are $l = 20d \times 20d \times 120d$ for simulations of designs with $L_{\max} = 20$, and $l = 24.5d \times 24.5d \times 120d$ for simulations of designs with $L_{\max} = 30$. Each direct-coexistence simulation contains a total of $N_{\text{tot}} = 888$ chains for the four-component case and $N_{\text{tot}} = 900$ chains for the six-component cases. We perform NVT molecular dynamics simulations using the Nose–Hoover thermostat and a timestep of $5 \times 10^{-3}\tau$ (in LJ units).

To verify equilibration, we calculate the degree of mixing (d.o.m.) over the region of the simulation box that is occupied by condensed phases (Fig. S1). The d.o.m. is defined to be

$$\text{d.o.m.} \equiv 1 - \frac{1}{2} \int_{\phi_{\text{T}}(z) > 1/2} dz \, \frac{\left[p_{\text{init}}^{(\beta)}(z) - p_{\text{init}}^{(\alpha)}(z)\right]^2}{p_{\text{init}}^{(\alpha)}(z) + p_{\text{init}}^{(\beta)}(z)}, \tag{S23}$$

where $p_{\text{init}}^{(\alpha)}(z)$ and $p_{\text{init}}^{(\beta)}(z)$ represent the probability of finding a chain that was initialized in the $\alpha$ or $\beta$ condensed phase, respectively, at the position $z$ along the simulation box. These probabilities are normalized such that $\int_{\phi_{\text{T}}(z) > 1/2} dz \, p_{\text{init}}^{(\alpha)} = 1$. If the chains completely mix, such that there is no correlation between the location of a particular chain and the condensed phase in which it was initially placed, then the degree of mixing tends to one. If the chains remain in their original phases, then the degree of mixing is close to zero. In situations where one or more species are shared between a pair of condensed phases, then the degree of mixing is expected to plateau at a value between zero and one. In practice, we use the degree of mixing to determine the timescale over which equilibration takes place in $\alpha = \beta$ control simulations (Fig. S1A). We also verify that the degree of mixing for $\alpha \neq \beta$ direct-coexistence simulations reaches a plateau value in the course of a simulation trajectory that is at least twice as long as the control-simulation mixing timescale (Fig. S1B–C).

#### D. Defining the heteropolymer design "fitness" metric

We compare alternative polymer designs by introducing an intuitive fitness metric based on the target order parameter profiles,

$$\text{fitness} \equiv \frac{\int_{\phi_{\rm T}(z)>1/2} dz\, \max_\alpha \left[q^{(\alpha)}(\vec{\phi}(z))\right]}{\int_{\phi_{\rm T}(z)>1/2} dz}, \tag{S24}$$

where the order parameter is defined to be

$$q^{(\alpha)}(\vec{\phi}) \equiv \frac{\vec{\phi}\cdot\vec{\phi}^{(\alpha)}}{||\vec{\phi}||\,||\vec{\phi}^{(\alpha)}||}. \tag{S25}$$

The fitness metric is equal to one if the bulk molecular concentrations in the condensed phases are precisely equal to the target molecular concentrations and the interfaces are perfectly sharp. However, in direct-coexistence simulations with $\alpha$, $\beta$, and dilute phases, a valid design will lead to (at least) three finite-width interfaces. A lower bound on the fitness of a valid design can be estimated by assuming that the integrand in Eq. (S24) is zero in the interfacial regions and that there are three interfacial regions, each of which comprises at most 5% of the total region in which $\phi_{\rm T} \geq 1/2$ (based on the typical interfacial width in the control simulations shown in Fig. S1A). In this way, we estimate that a valid design should have a fitness score of at least 0.85. We use this value as the threshold for determining whether a heteropolymer design is a success or a failure in Figs. 3 and 5 in the main text. We note that this heuristic fitness metric might not work as well as the number of components increases, since the order parameter is based on a Euclidean distance, and it generally becomes harder to distinguish points in high-dimensional spaces in this way due to the curse of dimensionality.

The equilibrated concentration and order-parameter profiles of the alternative designs for the example four-component design problem (see Fig. 3 in the main text) are shown in Fig. S2.

#### E. Computing effective polymer–polymer interactions in dilute and condensed phases

##### 1. *Effective polymer–polymer interactions in the dilute limit*

Polymer–polymer interactions in the dilute limit can be quantified by second virial coefficients [14],

$$B_{ij}^{\rm sim} = 2\pi \int_0^\infty dr\, r^2 \left\{1 - \exp\left[-w_{ij}(r)/k_{\rm B}T\right]\right\}, \tag{S26}$$

where $w_{ij}(r)$ is the simulated potential of mean force (PMF) between the centers of mass of two polymers of types $i$ and $j$. In practice, we compute $w_{ij}(r)$ using adaptive biasing force (ABF) simulations [15] implemented via the COLVARS package [16] in LAMMPS [11]. We run constant-temperature-and-volume (NVT) simulations with two chains initialized in a $20d \times 20d \times 20d$ simulation box with shrink-wrapped boundary conditions. Force statistics are stored in bins of width $0.25d$. The biasing force is applied once 1000 samples are collected in each bin, after which a final production run is performed for $10^8$ steps with a timestep of $0.005\tau$ (in LJ units). The second virial coefficient is then obtained by integrating the PMF using Eq. (S26). In Fig. 4a in the main text, five independent ABF simulations are performed for each $B_{ij}$ calculation to determine the statistical errors, shown as error bars in Fig. 4A in the main text.

We then compare the second virial coefficients computed via simulation to those predicted by the Flory–Huggins model,

$$B_{ij}^{\rm FH} = \frac{d^3 L_i L_j (1 + \epsilon_{ij}/k_{\rm B}T)}{2}. \tag{S27}$$

This comparison is shown in Fig. 4A in the main text.

##### 2. *Effective polymer–polymer interactions in condensed phases: Excess chemical potential differences*

We extract the excess chemical potential differences, $\Delta_{\alpha\beta}\mu_{{\rm ex},i} \equiv \mu_{{\rm ex},i}^{(\beta)} - \mu_{{\rm ex},i}^{(\alpha)}$, from the equilibrium compositions, determined from direct-coexistence simulations, of simulated $\alpha$ and $\beta$ bulk phases via [17]

$$\Delta_{\alpha\beta}\mu_{{\rm ex},i}^{\rm sim} = k_{\rm B}T \log\left(\langle\phi_i^{(\alpha)}\rangle / \langle\phi_i^{(\beta)}\rangle\right). \tag{S28}$$

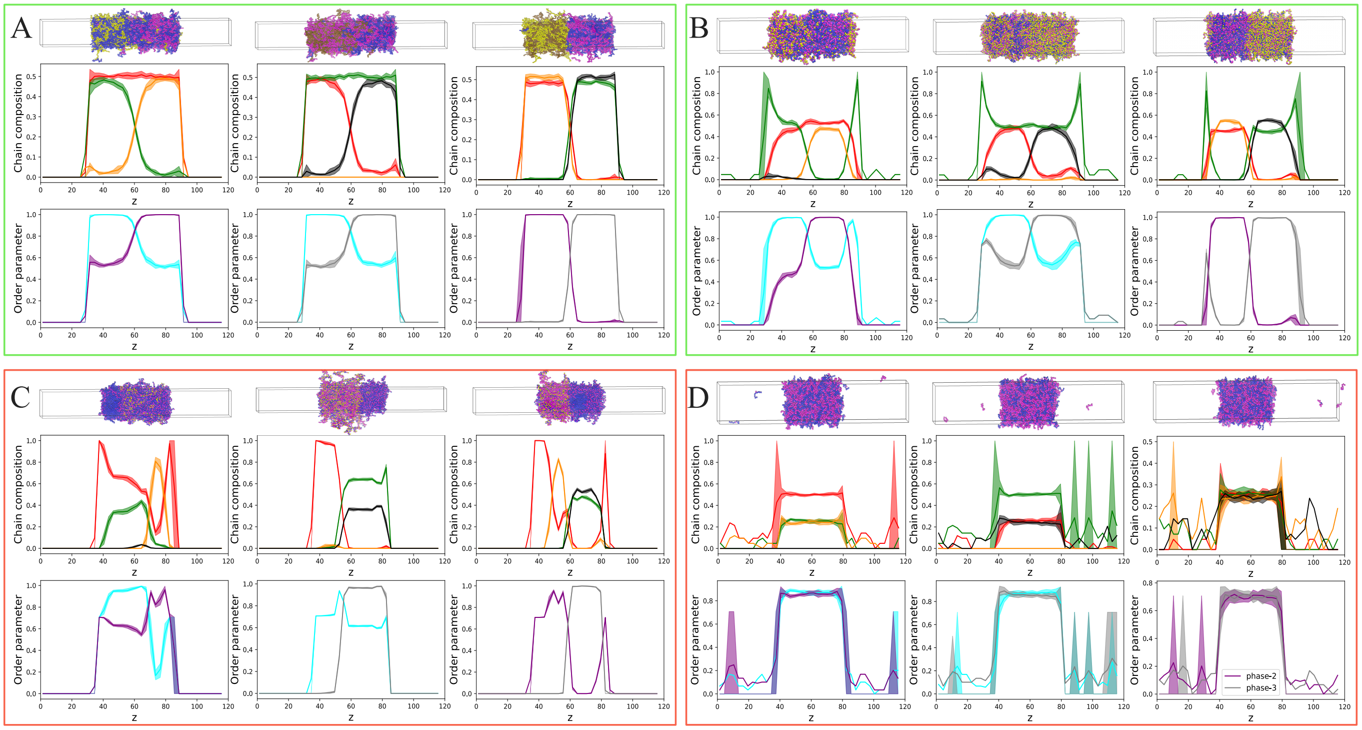


FIG. S2. **Simulation results for alternative design strategies**. Simulation results are shown for the alternative LJ heteropolymer designs summarized in Fig. 3b of the main text. Data are presented as in Fig. 3a of the main text. (A) A full-rank $r = 4$ design. The interaction matrix is obtained by minimizing the variance of the independent entries of $\boldsymbol{\epsilon}$, as in Ref. [2]. (B) The $r = 3$ design shown in Fig. 3a in the main text. (C) A design obtained by running the NMF solver on the interaction matrix used in **A**, but enforcing $r = 3$ monomer types. (D) A design obtained by running the NMF solver on the optimized $r = 3$ interaction matrix used in **B**, but enforcing $r = 2$ monomer types. The green and red outlines around each panel indicate whether the design is determined to be a success or a failure, respectively, according to the fitness metric, Eq. (S24).

We then compare these calculations with the Flory–Huggins prediction,

$$\Delta_{\alpha\beta}\mu_{\mathrm{ex},i}^{\mathrm{FH}} = -k_{\mathrm{B}}T\log[(1-\phi_{\mathrm{T}}^{(\beta)})/(1-\phi_{\mathrm{T}}^{(\alpha)})] + \sum_j \epsilon_{ij}(\phi_j^{(\beta)} - \phi_j^{(\alpha)}). \tag{S29}$$

This comparison is shown in Fig. 4B in the main text.

#### 3. *Effective polymer–polymer interactions in condensed phases: Radial distribution functions and structure factors*

For each condensed phase, we run an NPT simulation at zero pressure to compute the radial distribution function (RDF) with respect to the center of mass distance between polymers of various species (Fig. S3A). The RDF between chain types $i$ and $j$ is denoted $g_{ij}(r)$. Note that this calculation can only be performed with sufficient statistical accuracy for chain types that are enriched in a particular condensed phase. From the RDFs, we measure the well depths of the interactions, $\min_r -\log g_{ij}(r)$, which strongly correlate with the predicted pairwise interactions, $\epsilon_{ij}$ (Fig. S3B). This correspondence provides additional support for our conclusion that the mean-field approximations hold better in the condensed phases than in the dilute phase.

The mean-field interactions can also be related to the RDFs using linear response theory [14]. Specifically, the partial structure factor is related to the Fourier transform of the pair correlation function, $\hat{h} \equiv \hat{g} - 1$, via

$$S_{ij}(\vec{k}) = x_i\delta_{ij} + x_i x_j \rho \hat{h}_{ij}(\vec{k}), \tag{S30}$$

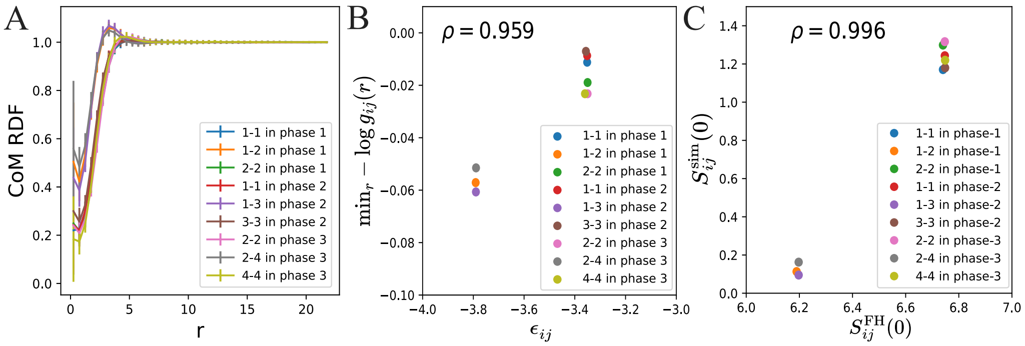


FIG. S3. **Effective polymer–polymer interactions determined from radial distribution functions (RDFs).** (A) RDFs, $g_{ij}(r)$, between the centers of mass (CoMs) of chains in the condensed phases. Only enriched chain types are shown for each condensed phase. (B) Correlations between the effective well depths inferred from the RDFs (see text) and the pairwise interactions, $\epsilon_{ij}$. The legend indicates pairs of chain types. (C) Correlations between the zero-mode partial structure factors, $S_{ij}(0)$, determined from the RDFs and the mean-field interactions in the Flory–Huggins model. In **B** and **C**, the Pearson correlation coefficient, $\rho$, is indicated.

where $x_i = \phi_i/\phi_\mathrm{T}$ is the chain composition in a condensed phase, $\rho = N_\mathrm{tot}/V$ is the total number density in the condensed phase, and $V$ is the simulated volume of the condensed phase. It follows that

$$\begin{aligned}\lim_{|\vec{k}|\to 0} S_{ij}(\vec{k}) &= x_i\delta_{ij} + 2x_i x_j \rho \int_0^\infty dr\, r h(r) \times \lim_{|\vec{k}|\to 0} \frac{\sin(2\pi r|\vec{k}|)}{|\vec{k}|} \\ &= x_i\delta_{ij} + 4\pi x_i x_j \rho \int_0^\infty dr\, r^2 h(r).\end{aligned} \tag{S31}$$

Finally, from linear response theory [14], we find

$$S_{ij}(0) = V^{-1}\frac{\partial\langle N_i\rangle}{\partial\beta\mu_j} = k_\mathrm{B}T\left(\frac{\partial\mu_j}{\partial\rho_i}\right)^{-1} = k_\mathrm{B}T\left(L_i\frac{\partial\mu_j}{\partial\phi_i}\right)^{-1}. \tag{S32}$$

We calculate the zero mode for number density fluctuations, $S_{ij}(0)$, via the mean-field expression, Eq. (S20)d, and the simulated single-phase RDFs, Eq. (S31), and find that the results are highly correlated (Fig. S3C). This finding also supports our conclusion that the mean-field approximations hold well in the condensed phases.

We compute the overlap parameter, $P$ [10], from these single-phase simulations by estimating the median number of distinct neighboring chains with which every chain in a condensed phase interacts. A chain is considered to be an interacting neighbor if there is at least one inter-chain pair of monomers that are within the cutoff distance, $3d$, of the monomer–monomer LJ potential.

### IV. DESIGN AND SIMULATION DATA FOR 6-COMPONENT HETEROPOLYMER DESIGNS

In this section, we present the optimization results and resulting simulation data for the problems considered in Figs. 5A and 5C in the main text. The results of convex optimization for both problems are shown in Fig. S4 using the same format as in Fig. 3A of the main text. For the condensate design problem shown in Fig. 5A, we select the optimal solution using the parameters $\lambda_\mathrm{min} = 0.5k_\mathrm{B}T$ and $L_\mathrm{max} = 20$. For the condensate design problem shown in Fig. 5C, we select the optimal solution using the parameters $\lambda_\mathrm{min} = 0.075k_\mathrm{B}T$ and $L_\mathrm{max} = 30$.

The LJ coefficients used for simulating the heteropolymer design shown in Fig. 5A in the main text are

$$\boldsymbol{w}^\mathrm{LJ} = \begin{bmatrix} 0.630 & 0.559 & 0.430 & 0.264 \\ 0.559 & 0.582 & 0.374 & 0.877 \\ 0.430 & 0.374 & 0.516 & 0.361 \\ 0.264 & 0.877 & 0.361 & 0.663 \end{bmatrix}.$$

The LJ coefficients used for simulating the heteropolymer design shown in Fig. 5C in the main text are

$$\boldsymbol{w}^\mathrm{LJ} = \begin{bmatrix} 0.135 & 0.701 & 0.482 & 0.457 \\ 0.701 & 0.483 & 0.271 & 0.375 \\ 0.482 & 0.271 & 0.951 & 0.525 \\ 0.457 & 0.375 & 0.525 & 0.982 \end{bmatrix}.$$

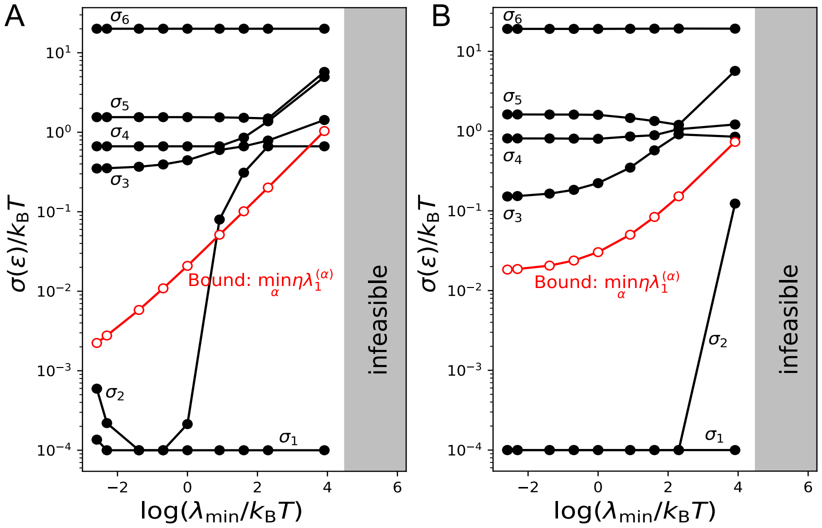


FIG. S4. **Convex optimization results for example six-component condensate design problems.** Data are shown as in Fig. 3A of the main text. (A) Convex optimization results for the condensate design problem shown in Fig. 5A in the main text. (B) Convex optimization results for the condensate design problem shown in Fig. 5C in the main text.

The corresponding simulation data for these designs, along with the optimized sequences shown in Figs. 5A and 5C in the main text, are shown in Fig. S5.

[1] C. Eckart and G. Young, Psychometrika **1**, 211–218 (1936).
[2] F. Chen and W. M. Jacobs, J. Chem. Phys. **158**, 214118 (2023).
[3] S. P. Boyd and L. Vandenberghe, *Convex optimization* (Cambridge University Press, 2004).
[4] S. Diamond and S. Boyd, J. Mach. Learn. Res. **17**, 2909 (2016).
[5] B. O'Donoghue, E. Chu, N. Parikh, and S. Boyd, J. Optimiz. Theory App. **169**, 1042 (2016).
[6] B. Recht, M. Fazel, and P. A. Parrilo, SIAM Review **52**, 471–501 (2010).
[7] Z. Yang and E. Oja, Pattern Recognition **45**, 1500 (2012).
[8] S. C. Glotzer and M. J. Solomon, Nat. Mater. **6**, 557 (2007).
[9] P.-T. De Boer, D. P. Kroese, S. Mannor, and R. Y. Rubinstein, Ann. Oper. Res. **134**, 19 (2005).
[10] R. H. Colby and M. Rubinstein, *Polymer physics* (Oxford University Press, 2003).
[11] S. Plimpton, J. Comput. Phys. **117**, 1 (1995).
[12] J. E. Lennard-Jones, Proc. Royal Soc. A **106**, 463–477 (1924).
[13] K. Kremer and G. S. Grest, J. Chem. Phys. **92**, 5057–5086 (1990).
[14] J.-P. Hansen and I. R. McDonald, *Theory of simple liquids: With applications to soft matter* (Academic Press, 2013).
[15] E. Darve, D. Rodríguez-Gómez, and A. Pohorille, J. Chem. Phys. **128** (2008).
[16] J. Hénin, G. Fiorin, C. Chipot, and M. L. Klein, Journal of Chemical Theory and Computation **6**, 35–47 (2009).
[17] W. M. Jacobs, J. Chem. Theory Comput. **19**, 3429 (2023).

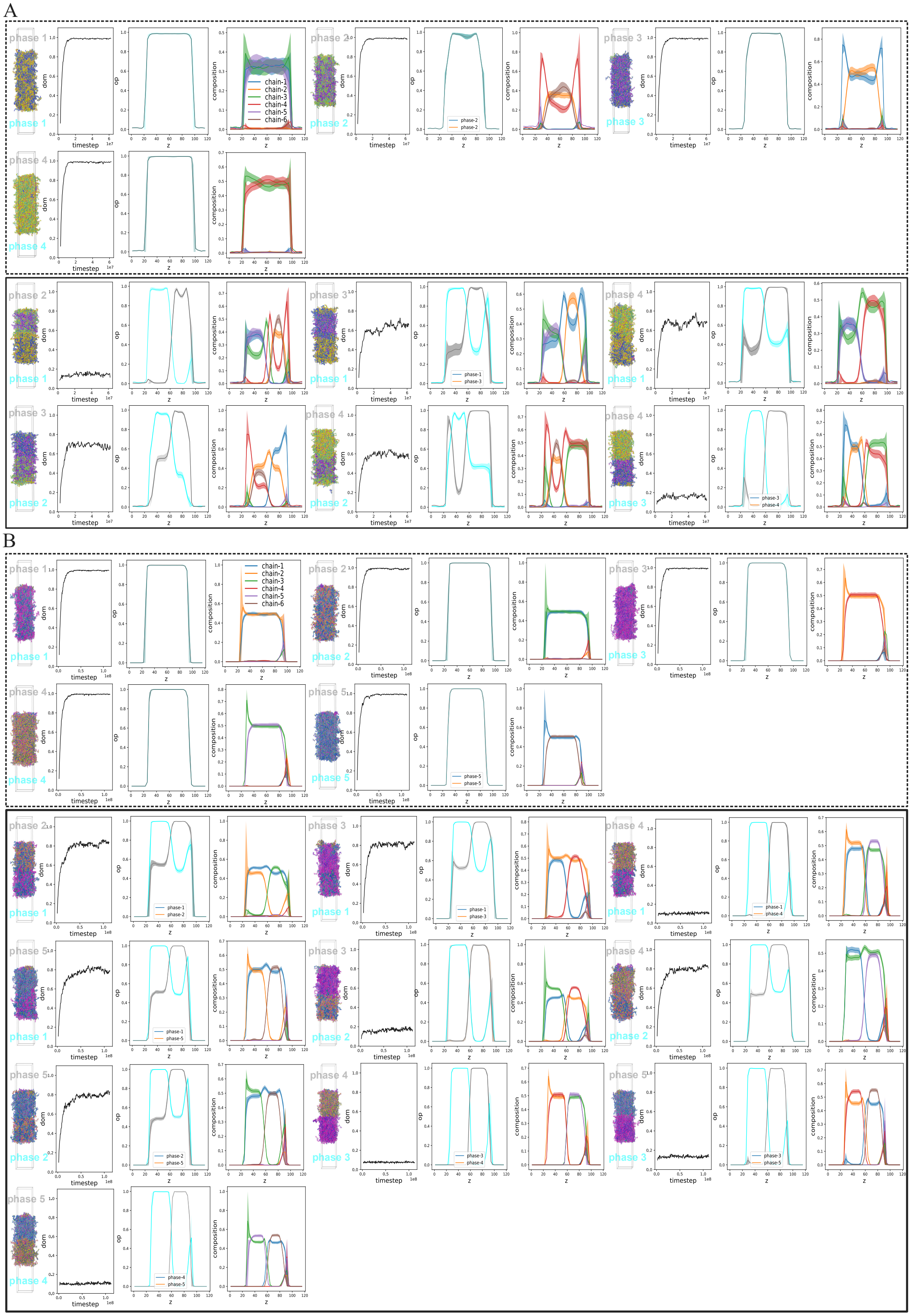


FIG. S5. **Molecular simulation results for optimized LJ heteropolymer solutions to example six-component condensate design problems.** Simulation results of all pairs of coexisting phases are shown for (A) the condensate design problem presented in Fig. 5A of the main text and (B) Fig. 5C of the main text. Results in dashed boxes correspond to control simulations, for which $\alpha = \beta$. Results in solid boxes correspond to direct-coexistence simulations of immiscible phases, for which $\alpha \neq \beta$.
